# Distal mutations in a designed retro-aldolase alter loop dynamics to shift and accelerate the rate-limiting step

**DOI:** 10.1101/2025.01.26.634918

**Authors:** Serena E. Hunt, Cindy Klaus, Aqza E. John, Niayesh Zarifi, Alec Martinez, Ferran Feixas, Marc Garcia-Borràs, Michael C. Thompson, Roberto A. Chica

## Abstract

Amino-acid residues distant from an enzyme’s active site are known to influence catalysis, but their mechanistic contributions to the catalytic cycle remain poorly understood. Here, we investigate the structural, functional, and mechanistic impacts of distal and active-site mutations discovered through directed evolution of the computationally designed retro-aldolase RA95. Active-site mutations improve catalytic efficiency by 3,600-fold, while distal mutations alone offer no improvement. When combined with active-site mutations, distal mutations further increase efficiency by 6-fold, demonstrating an epistatic effect. X-ray crystallography and molecular dynamics simulations reveal that distal mutations promote active site opening by altering loop dynamics. Kinetic solvent viscosity effects and electrostatic analysis show that distal mutations accelerate the chemical transformation by 100-fold, shifting the rate-limiting step to product release, which is further accelerated by the increased opening of the active site. These findings highlight the critical role of distal residues in shaping the active-site environment and facilitating the structural dynamics essential for progression through the catalytic cycle.

## Introduction

Enzymes accelerate chemical reactions by many orders of magnitude, enabling life to operate within biologically relevant timescales. Although decades of biochemical and structural studies have provided deep insights into the role of active-site residues in catalysis (*1–3*), the contribution of distal regions in promoting the catalytic cycle via allosteric interactions remains poorly understood (*4, 5*). This knowledge gap hinders our ability to predict the effects of distal mutations on enzyme function, limiting our understanding of disease-causing mutations and preventing the design of proficient artificial enzymes. Recent molecular dynamics studies on the effects of distal mutations in enzymes improved through directed evolution suggest that these mutations contribute to alter networks of non-covalent interactions, redistributing conformational states within the ensemble to favor productive ones (*6, 7*). These changes often involve flexible loops and lids that regulate access to the active site or shape the binding site to modulate substrate binding and active-site preorganization (*8–12*). However, previous studies have investigated the role of distal mutations alongside active-site mutations, making it difficult to determine whether their effects on catalysis are direct or arise from epistatic interactions with active-site mutations. Furthermore, the mechanistic effects of distal mutations on the catalytic cycle have been largely overlooked, preventing a full understanding of how these mutations impact various steps along the reaction coordinate and contribute to overall catalytic efficiency.

The de novo retro-aldolase RA95 (*13*) offers a compelling model for understanding the role of distal mutations in facilitating the catalytic cycle. RA95 was initially designed to catalyze the retro-aldol decomposition of methodol (Figure 1a) by sculpting an active site for this reaction within a natural protein scaffold that lacks this function. Its initial catalytic activity was modest (*k*_cat_ = 5 × 10^−5^ s^−1^), but directed evolution improved this by five orders of magnitude through 19 rounds that introduced a total of 22 mutations, yielding the final evolved variant RA95.5-8F (*14, 15*) (Figure 1b). Unlike other de novo enzymes subjected to directed evolution (*16–22*), the evolution of RA95 involved substantial active-site remodeling. This included replacement of the original catalytic nucleophile (Lys210) with a new one (Lys83) and introduction of three additional residues (Tyr51, Asn110 and Tyr180) to form a catalytic tetrad that enhances catalysis through a hydrogen bond network (*15*) (Figure 1c). Mutations also triggered conformational shifts in nearby surface loops to relieve steric clashes with the new substrate binding position in the active site (Figure 1d). While the effects of these active site changes can be rationalized, the role of distal mutations in facilitating these structural adjustments and accelerating the catalytic cycle remains unclear.

**Figure 1.**
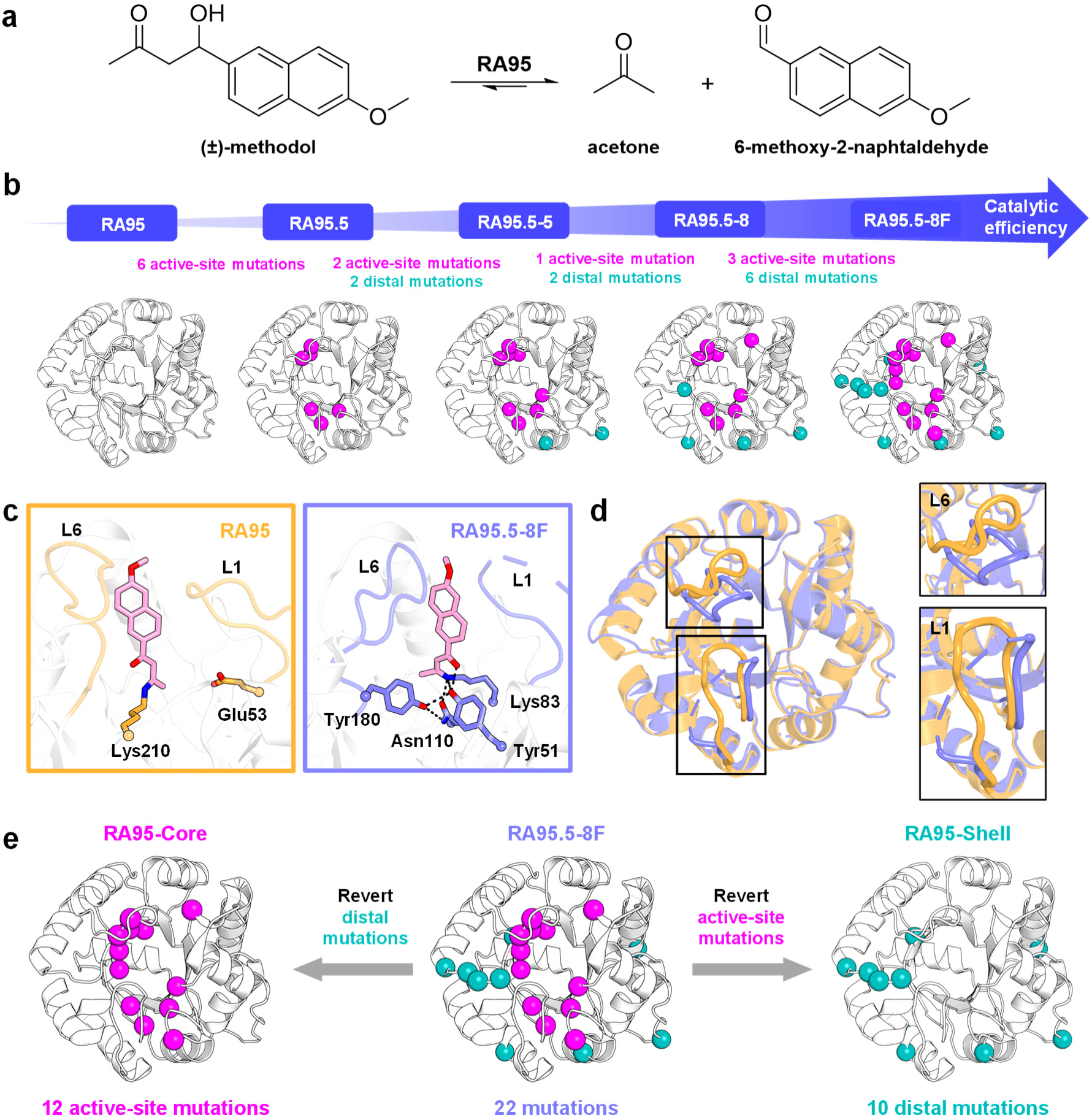
RA95 series of retro-aldolases. (a) Retro-aldolases catalyze the multi-step carbon–carbon bond cleavage of 4-hydroxy-4-(6-methoxy-2-naphthyl)-2-butanone (methodol) into 6-methoxy-2-naphthaldehyde and acetone. (b) Evolutionary trajectory of the computationally designed de novo retro-aldolase RA95 spanning the variants RA95.5, RA95.5-5, RA95.5-8, and RA95.5-8F. A combination of 12 active site mutations (magenta spheres) and 10 distal mutations (teal spheres) were introduced over 19 rounds of directed evolution. If a position was mutated multiple times along the evolutionary trajectory, the mutation is only shown in the variant where it was mutated for the last time. (c) Active sites of RA95 (orange, PDB ID: 4A29) (*14*) and RA95.5-8F (purple, PDB ID: 5AN7) (*15*) show catalytic residues and the covalent diketone inhibitor used in crystallization (pink). Active-site loops L1 (residues 52– 66) and L6 (residues 180–190) are indicated. The catalytic motif that was designed in RA95 comprises a nucleophilic lysine (Lys210) and a glutamate (Glu53) positioned nearby to orient a catalytic water molecule. Through evolution, a tetrad comprising a new catalytic lysine (Lys83) and three additional residues participating in a hydrogen bond network (Tyr51, Asn110, and Tyr180) was created. (d) Directed evolution resulted in conformational changes to loops L1 and L6 (rectangles). In RA95.5-8F, residues 58–63 of loop L1 are disordered, resulting in a gap in the electron density indicated by a dashed line. The structures of RA95 and RA95.5-8F are shown in orange and purple, respectively. (e) RA95-Core and RA95-Shell are variants of RA95 that contain either active-site or distal mutations identified by directed evolution of RA95.5-8F.

In this study, we investigate how distal mutations introduced through directed evolution promote the RA95 catalytic cycle and enhance its overall efficiency. Our findings show that distal mutations augment activity by shifting the rate-limiting step to product release and accelerating this process. This enhancement is driven by altered surface loop motions that facilitate active site opening while simultaneously optimizing the local electric field at the active site. These results underscore the multifaceted role of distal mutations in modulating loop dynamics, enhancing transition-state stabilization, and facilitating product release, ultimately lowering energy barriers across multiple steps to accelerate the catalytic cycle. The mechanistic insights reported here illuminate the ways in which distal mutations can enhance catalytic activity in the context of natural evolution and de novo enzyme engineering, or disrupt catalytic activity in the context of human disease mutations.

## Results

### Functional effects

To investigate the functional effects of distal mutations introduced during directed evolution of RA95, we created two enzyme variants in which either the distal or active-site mutations from the final evolved variant, RA95.5-8F, were reverted to their identities in the original designed enzyme. We call these variants RA95-Core and RA95-Shell, respectively (Figure 1e, Supplementary Table 1). We define active-site mutations as those found within 8 Å of the diketone inhibitor that forms a covalent bond with the catalytic Lys83 residue in the crystal structure of RA95.5-8F (PDB ID: 5AN7) (*15*). Mutations occurring at residues beyond this 8-Å radius are considered distal. We chose the 8 Å cutoff to include all residues in direct contact with the inhibitor (first shell) and those interacting with the first-shell residues (second shell), which are typically targeted in de novo enzyme design (*13, 23–25*). Given the different active-site configurations and catalytic motifs between RA95 and RA95.5-8F (Figure 1c), we postulated that analyzing these enzymes alongside Core and Shell variants would yield insights on the catalytic role of distal mutations that may have been obscured in previous analyses of RA95 directed evolution variants (*6, 14, 15, 26*) containing both active-site and distal mutations (Figure 1b, Supplementary Table 1).

Kinetic characterization of RA95-Core revealed that it catalyzes the cleavage of (±)- methodol with a catalytic efficiency of 1,900 M^−1^ s^−1^, a 3,600-fold increase compared to RA95 (Table 1, Supplementary Figure 1). This efficiency is higher that that of the penultimate evolutionary intermediate, RA95.5-8 (*k*_cat_/*K*_M_ = 850 M^−1^ s^−1^) (*14*), but 6-fold lower than RA95.5-8F. Given that the pKa of the RA95-Core catalytic lysine is within error to that of RA95.5-8F (Table 1, Supplementary Figure 2), these results indicate that the 6-fold lower activity of RA95-Core is not due to a pKa difference affecting the nucleophilicity of the catalytic lysine.

**Table 1.** Kinetic parameters of retro-aldolase variants.

| Enzyme | $k_{\text{cat}}$<br>(s<sup>-1</sup>)<sup>a</sup> | $K_M$<br>(μM<sup>-1</sup>)<sup>a</sup> | $k_{\text{cat}}/K_M$<br>(M<sup>-1</sup> s<sup>-1</sup>) | pK<sub>a</sub><sup>b</sup> | T<sub>m</sub><sup>c</sup><br>(°C) |
| --- | --- | --- | --- | --- | --- |
| RA95 | 0.00027 ± 0.00004 | 500 ± 200 | 0.52 | 8.1<sup>d</sup> | 83.66 ± 0.02 |
| RA95.5-8F | 4.6 ± 0.3 | 390 ± 50 | 12,000 | 5.6 ± 0.1 | 71.9 ± 0.1 |
| RA95-Shell | 0.00016 ± 0.00002 | 400 ± 100 | 0.37 | - | 81.56 ± 0.03 |
| RA95-Core | 0.32 ± 0.01 | 170 ± 10 | 1,900 | 5.8 ± 0.1 | 68.7 ± 0.1 |
| RA95-Core-Y51F | 0.026 ± 0.001 | 240 ± 20 | 110 | 5.99 ± 0.08 | - |
| RA95-Core-N110S | 0.114 ± 0.003 | 130 ± 10 | 880 | 5.63 ± 0.07 | - |
| RA95-Core-Y180F | 0.05 ± 0.01 | 170 ± 30 | 290 | 6.53 ± 0.06 | - |

<sup>a</sup> Kinetic parameters were determined for (±)-methodol. $k_{\text{cat}}$ and $K_M$ were calculated by fitting the data to the Michaelis-Menten model $v_0 =$ $k_{\text{cat}}[E_0][S]/(K_M + [S])$ . Errors of nonlinear regression fitting are provided. $n =$ six or nine individual replicates performed on two or three independent enzyme batches.
<sup>b</sup> $k_{\text{cat}}/K_M$ versus pH data were fitted to the following equation using nonlinear least squares regression: $(k_{\text{cat}}/K_M)_{\text{obs}} = (k_{\text{cat}}/K_M)_{\text{max}}/(1 + 10^{\text{pK}_{a1} - \text{pH}} +$ $10^{\text{pK}_{a2} - \text{pH}})$ . The apparent pK<sub>a</sub> of the catalytic lysine (pK<sub>a1</sub>) of each variant is presented, with the errors of nonlinear regression fitting provided.
<sup>c</sup> Thermal denaturation midpoint temperature determined through loss of CD signal at 222 nm. Errors of nonlinear regression fitting to a two-state transition model are provided.
<sup>d</sup> Value from (14).

Furthermore, each of the catalytic tetrad residues in RA95-Core contributes to enhanced catalysis as mutation of these residues to disrupt hydrogen-bonding interactions results in 3–12-fold decreases in *k*_cat_ (Table 1), with the Tyr51Phe mutation having the biggest impact. These results align with the trend seen when equivalent mutations were introduced into RA95.5-8F (*15*). Together with the pKa measurements, these mutational studies suggest that the active-site configuration of RA95-Core is similar to that of the evolved variant, featuring an identical catalytic tetrad.

By contrast, kinetic characterization of RA95-Shell showed that distal mutations alone decrease *k*_cat_ by almost two-fold (Table 1, Supplementary Figure 1). However, when active-site mutations are introduced into RA95-Shell to form the evolved variant, they result in a 29,000-fold increase in *k*_cat_, demonstrating synergistic effects specific to the evolved active site. Synergistic effects between distal and active-site mutations are also observed in the thermal stability of retro-aldolase variants (Table 1, Supplementary Figure 3). For example, adding distal mutations to RA95 lowers its melting temperature by approximately 2 °C, whereas active-site mutations are highly destabilizing, reducing the melting temperature by 15 °C. However, when distal mutations are introduced into RA95-Core to form RA95.5-8F, the melting temperature increases by approximately 3 °C. These results suggest that distal mutations were selected by evolution not only for their beneficial impact on catalytic activity but also to partially compensate for the large destabilization caused by the optimized active-site configuration of RA95.5-8F.

### Structural effects

To investigate how changes to the enzyme structure caused the observed activity effects, we turned to X-ray crystallography. We targeted structures of our enzymes in their unbound form to assess the structural impact of mutations on the RA95 fold without potential rearrangements caused by ligand binding (Figure 1c,d). We successfully grew crystals for RA95-Shell but were unable to do so for RA95-Core. Additionally, we crystallized RA95, as its structure without a covalent inhibitor was not previously available. The unit cells for RA95-Shell and RA95 corresponded to space group P 21 21 2 with one protein molecule in the asymmetric unit, and they diffracted at resolutions of 1.77 Å and 1.89 Å, respectively (Supplementary Table 2). Comparison of these crystal structures with the previously published structure of RA95.5-8F in its unbound form (PDB ID: 5AOU (*15*)) revealed conformational changes in active-site loops L1 (residues 52– 66) and L6 (residues 180–190). In RA95, loop L6 adopts a conformation that positions it further away from loop L1 than in RA95.5-8F (Figure 2a), and this distance increases to accommodate the bound inhibitor (PDB ID: 4A29 (*14*)) (Supplementary Figure 4a). By contrast, there is no substantial change in the conformation of loops L1 or L6 upon inhibitor binding in RA95.5-8F (Supplementary Figure 4b), suggesting that these loops are already positioned for efficient substrate binding. However, both the bound and unbound structures of the evolved enzyme show no density for residues 58–61 and 58–63 of loop L1, respectively, indicating that one side of this loop is disordered. This result contrasts with RA95, where clear density is observed for loop L1 in both bound and unbound forms (Supplementary Figure 5). Thus, the combination of active-site and distal mutations introduced by directed evolution remodelled surface loops in RA95.5-8F to enhance substrate recognition while also increasing conformational heterogeneity of loop L1.

**Figure 2.**
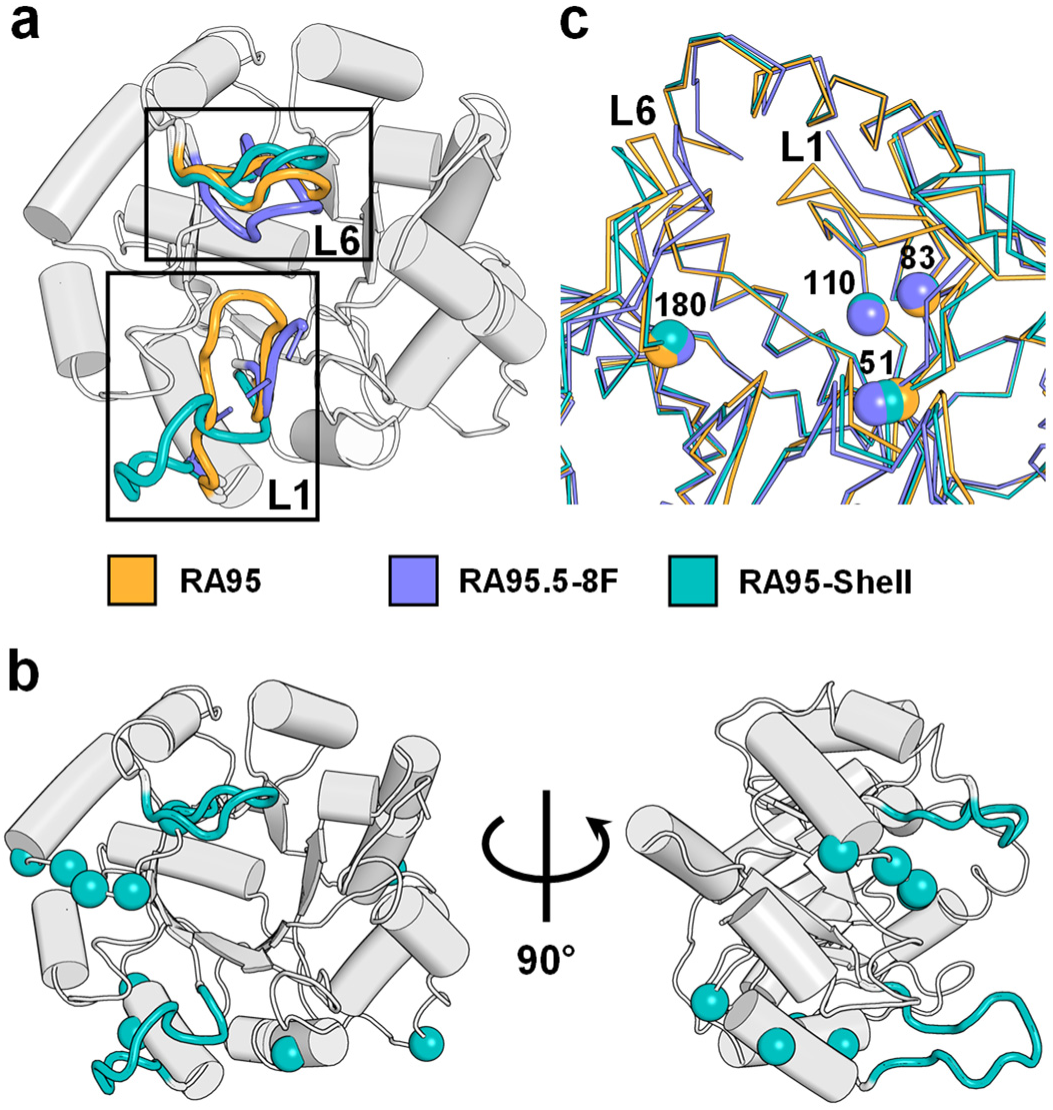
Structural effects of distal mutations. (a) Superposition of crystal structures for unbound RA95 (orange, PDB ID: 9MYA), RA95-Shell (teal, PDB ID: 9MYB) and RA95.5-8F (purple, PDB ID: 5AOU) (*15*). Loops L1 and L6 are indicated in rectangles and coloured while a representative retro-aldolase structure in grey is shown for the remainder of the protein. The dashed line indicates a gap in the electron density for loop L1 residues 58–63 in RA95.5-8F. (b) Crystal structure for unbound RA95-Shell. Loops L1 and L6 are shown in teal. Distal mutations (spheres) are not located on or near loops L1 or L6. (c) Superposition of ribbon representations of the unbound RA95 (orange), RA95-Shell (teal) and RA95.5-8F (purple) active sites. Cα carbons of positions 51, 83, 110, and 180 are shown as spheres and loops L1 and L6 are indicated.

In RA95-Shell, there is a large conformational change in loop L1 that positions it approximately 10 Å away from its position in RA95 (Figure 2a, Supplementary Figure 5), a conformation that has not been observed in any other crystallized retro-aldolase variant to date, and that cannot be predicted by AlphaFold2 (Supplementary Figure 6). Interestingly, this large conformational shift is caused by distal mutations that are not located on or near loops L1 or L6 (Figure 2b). The large conformational change in loop L1 is accompanied by a shift in loop L6, which moves further away from the position it adopts in the unbound structures of RA95 or RA95.5-8F, making the active site of RA95-Shell more open than any of the other variants. These findings could explain why distal mutations alone are detrimental to activity when introduced into RA95, as they lead to a conformation that is more open and dissimilar to the reactive conformation observed in the inhibitor-bound form of RA95. However, when combined with active-site mutations, distal mutations enable loops L1 and L6 to adopt conformations conducive to efficient catalysis, as seen in the structures of RA95.5-8F. The increased conformational heterogeneity of loop L1 in RA95.5-8F is likely caused by distal mutations, since these mutations alone can induce a large conformational change in this loop. Furthermore, this heterogeneity is absent in RA95, which lacks these mutations.

In addition to causing large conformational changes in active-site loops, distal mutations also induce more subtle shifts in the backbone position of active-site residues, despite being far from the mutation sites (Figure 2c). Notably, the Cα carbon at position 51 shifts by 0.7 Å when comparing RA95 to RA95-Shell, which causes minimal changes to the rotameric configuration of active-site residues, except for catalytic residue Lys210, which is already very flexible (Supplementary Figure 7). Given that Tyr51 emerged early in the RA95 evolutionary trajectory, this backbone shift may help position this residue optimally for its catalytic role in RA95.5-8F. The Cα carbon at position 51 shifts an additional 0.7 Å in RA95.5-8F compared to RA95-Shell, leading to an overall shift of 1.4 Å due to both active-site and distal mutations. Overall, our findings suggest that distal mutations can induce both large and subtle structural changes, likely contributing to the activity enhancements and stability changes seen during directed evolution.

### Dynamical effects

Given that distal mutations cause a large conformational change to loop L1 in RA95-Shell and contribute to its high conformational heterogeneity in RA95.5-8F, we investigated the effects of distal mutations on structural dynamics using microsecond-timescale molecular dynamics simulations. Structural differences along the molecular dynamics trajectories were analyzed using principal component analysis (PCA). This analysis revealed population shifts in conformational states due to the different combinations of mutations (Figure 3). The greatest variation across the dataset was driven by loop L1 residues 59–62. In agreement with the crystal structures described earlier, this loop interconverts during molecular dynamics between open and closed conformational states, which are classified according to the Cα distance between residues 58 and 185 on loops L1 and L6, respectively. The first principal component distinguishes between snapshots with a closed L1 conformation, as seen in the unbound RA95 crystal structure (distances between L1 and L6 around 13 Å), and those with an open L1 conformation, as observed in RA95-Shell (distances around 23 Å).

**Figure 3.**
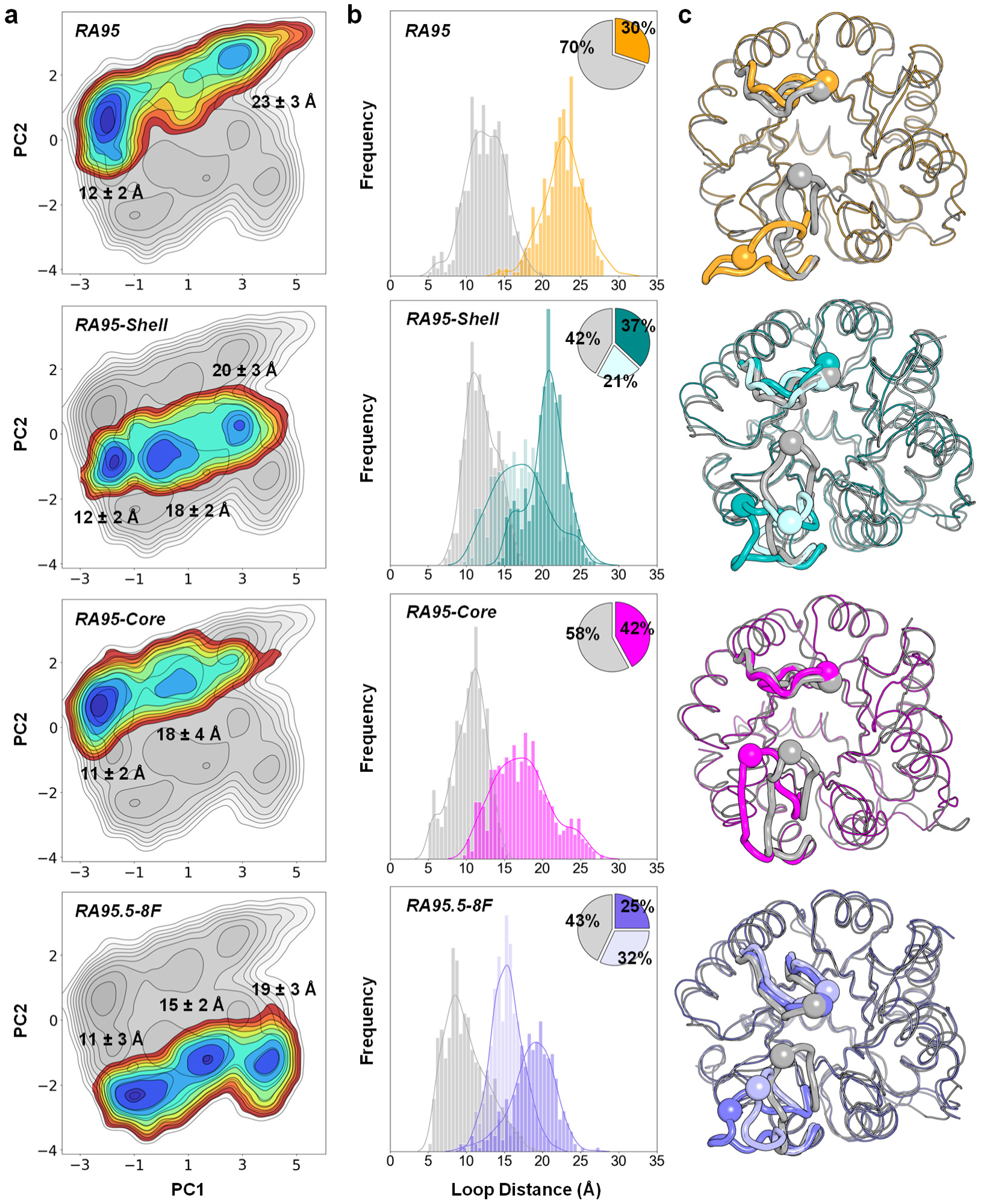
Dynamic effects of distal mutations. (a) Trajectories projected into the two most important principal components (PC1 and PC2) based on Cα contacts. Partitioning of the trajectories was performed using distance-based k-means clustering and the mean and standard deviation (in Å) of the distance between loops L1 and L6 is shown for each cluster. Loop distance describes the distance between the Cα carbons of residues 58 and 185. PC1 differentiates structures with closed active-site loops (low PC1 values, smaller loop distances) from those with open active site loops (higher PC1 values, larger loop distances). (b) Histograms of loop distances after partitioning of the trajectories. A pie chart showing the proportions of conformations in each cluster is shown for each variant. (c) Centroid structures of each cluster determined by computing pairwise root-mean-square deviations between all conformations of the cluster. Centroid structures are coloured according to their corresponding clusters in (b). The Cα carbons of residues 58 and 185 are shown as spheres. Active-site loops L1 and L6 are shown as thicker regions of the cartoon structure.

Comparison of the PCA plots for RA95 and RA95.5-8F (Figure 3a) shows that evolution alters the conformational landscape, shifting RA95 from two major conformational states (closed and open) to three distinct populations in RA95.5-8F (closed, partially open and open). This shift decreases the proportion of snapshots in the conformational ensemble where loop L1 adopts a closed conformation (Figure 3b,c) similar to the inhibitor-bound form of the enzyme. Notably, the increased prevalence of open and partially open conformations of loop L1 following evolution is attributed to the addition of distal mutations. Indeed, when distal mutations were introduced into RA95 to create RA95-Shell or RA95-Core to create RA95.5-8F, the conformational landscape shifts from two major states to three, which is accompanied by an increase in the proportion of snapshots where L1 is open or partially open (Figure 3b). Conversely, the addition of active-site mutations to RA95 (to form RA95-Core) or RA95-Shell (to create RA95.5-8F) reduces the proportion of open snapshots in the population towards closed or partially open snapshots. In RA95-Core, active-site mutations nearly eliminate the open conformation (loop distance of 23 ± 3 Å) and introduce a new state where L1 is partially open (loop distance of 18 ± 4 Å) (Figure 3a). These results demonstrate how distal mutations influence enzyme conformational dynamics, causing shifts in the conformational landscape that enrich open conformations and depopulate closed ones.

### Mechanistic effects

X-ray crystallography and molecular dynamics simulations showed that distal mutations favor opening of the active site, which could facilitate active-site accessibility. To investigate this possibility, we measured kinetic solvent viscosity effects on RA95-Core and RA95.5-8F using sucrose as the viscogen (Supplementary Figure 8). These experiments help determine if substrate binding is diffusion-controlled and if product release is the rate-limiting step in the catalytic cycle (*27*). In these analyses, substrate binding, product release and conformational changes in the enzyme structure are expected to be diffusion limited with rate constants dependent on the solvent viscosity. Conversely, the chemical step of catalysis is typically assumed to be independent of solvent viscosity because the chemical transformation itself occurs within the active site of the enzyme, where the environment is generally shielded from bulk solvent effects.

A plot of normalized *k*_cat_ as a function of relative solvent viscosity shows slopes between 0 and 1 for both RA95-Core and RA95.5-8F (Figure 4a), indicating that the overall turnover is partially limited by product release in both variants. From these slopes, rate constants for the chemical transformation (*k*_3_) and product release (*k*_4_) were calculated using equations 1 and 2 (Methods), revealing that distal mutations led to a 100-fold increase in *k*_3_ and a 4-fold increase in *k*_4_ (Figure 4b). These changes result in a shift in the rate-limiting step, from the chemical transformation in RA95-Core to product release in RA95.5-8F. Importantly, *k*_3_ and *k*_4_ values for RA95.5-8F are close to the previously reported values for single turnover C–C bond cleavage or earlier step (*k* = 35 ± 4 s^−1^) and enamine breakdown during acetone release (*k* = 5.2 ± 0.5 s^−1^), respectively (Figure 4b) (*26*). Since all steps in the proposed retro-aldolase mechanism leading to C–C bond cleavage are not expected to be affected by solvent viscosity (Figure 4c), we conclude that *k*_3_ corresponds to rate constant of the rate-limiting step on the C–C bond cleavage path. Previously, enamine degradation by acid protonation to form the Schiff base intermediate has been shown to be rate-limiting in RA95.5-8F (*26*). This step should be affected by viscosity since water has been proposed to be the acid that protonates this enamine, and water would need to diffuse into the active site to act as an acid. Furthermore, the Schiff base intermediate is hydrolyzed to produce acetone, whose release from the active site should also be affected by solvent viscosity. Thus, our results indicate that distal mutations enhance catalysis by facilitating product release (*k*_4_), which involves the rate-limiting enamine degradation leading to hydrolysis of the acetone/Lys adduct, likely through increased opening of the active site by altered dynamicity of loop L1.

**Figure 4.**
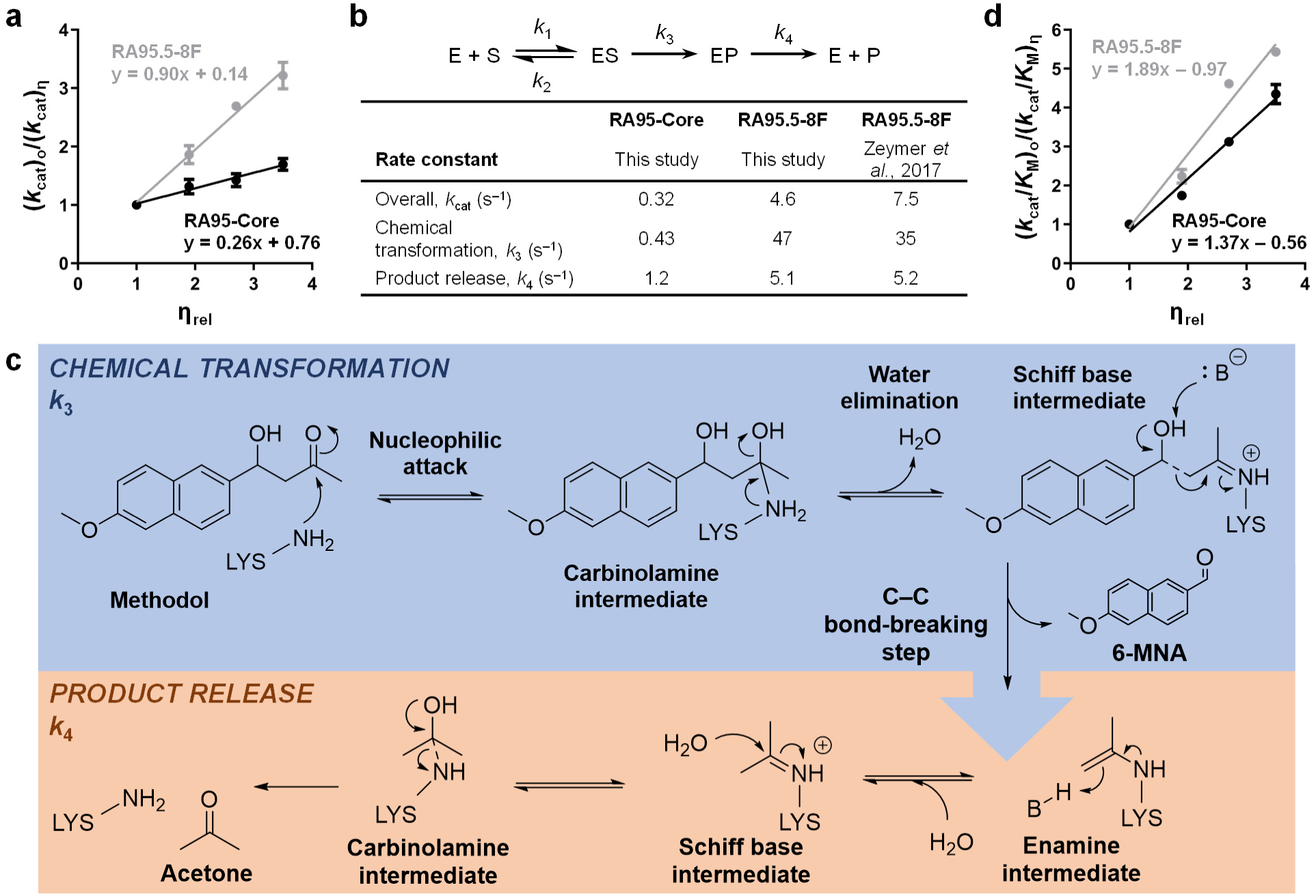
Mechanistic effects of distal mutations. (a) Kinetic solvent viscosity effects on *k*_cat_ provide insight into product release. Kinetic parameters, normalized to values obtained in a non-viscous buffer, are plotted against the relative viscosity (η_rel_). Data represent mean ± SEM from n = 2 independent biological replicates measured at various buffer viscosities. (b). Mechanism of an enzyme (E) reaction with a single substrate (S) and product (P), illustrating rate constants for substrate association (*k*_1_), chemical transformation (*k*_3_), and product release (*k*_5_). Rate constants were extracted from slopes of kinetic solvent viscosity effects and *k*_cat_ using equations 1 and 2 (Methods). Distal mutations resulted in a 100-fold increase in the rate of the chemical transformation (*k*_3_) and a 4-fold increase in the rate of product release (*k_5_*). (c) Based on the measured kinetic solvent viscosity effects, we propose that the chemical transformation step (*k*_3_) corresponds to the rate-limiting step of C–C bond cleavage, as this step is not expected to be influenced by solvent viscosity. By contrast, the product release step (*k*_4_) corresponds to the rate-limiting step of enamine degradation via acid protonation and Schiff-base hydrolysis to release acetone, both of which depend on solvent diffusion and are affected by viscosity. (d) Kinetic solvent viscosity effects on *k*_cat_/*K*_M_ provide insights into substrate capture in enzyme−substrate complexes that lead to product formation. Data represent the mean ± SEM for measurements from *n* = 2 independent biological replicates at various buffer viscosities.

By contrast, a plot of the normalized *k*_cat_/*K*_M_ as a function of relative solvent viscosity shows slopes greater than 1 for both variants (Figure 4d). Since a slope of 1 is the theoretical limit for diffusion-limited catalysis, slopes greater than 1 reflect inhibitory effects of the viscogen on the enzyme structure or additional diffusion-controlled equilibria not directly associated with substrate diffusion into the active site (*27*). Such slopes have previously been reported in enzymes where a viscosity dependent conformational change accompanies substrate binding (*28, 29*). Thus, our results suggest that a diffusion-limited conformational change accompanies substrate binding in both RA95-Core and RA95.5-8F. Greater viscosity effects are observed for RA95.5-8F, suggesting that these effects are caused by the increased conformational heterogeneity of loop L1 induced by distal mutations in RA95.5-8F. Taken together, these findings confirm that distal mutations contribute to catalysis in RA95.5-8F by shifting the rate-limiting step to product release and accelerating it through altered loop dynamics that increase active-site accessibility.

### Electrostatic effects

Distal mutations in RA95.5-8F introduce a net surface charge change of –4 by replacing three arginines with neutral amino acids and introducing a negatively charged aspartate residue (Supplementary Table 1). We hypothesized that this altered charge distribution modifies the local electric field (LEF) within the active site, potentially accounting for the observed 100-fold increase in *k*_3_, as LEF changes can affect reaction rates by modulating charge distribution, with previous studies showing up to 50-fold enhancements(*30–32*). To test this hypothesis, we analyzed the electrostatic preorganization(*33*) of RA95-Core and RA95.5-8F active sites by calculating the LEF at the catalytic center (Figure 5a) using molecular dynamics ensembles of open and closed conformational states. While LEF magnitudes were comparable across variants and conformational states, RA95.5-8F exhibited significantly different LEF orientations compared to RA95-Core, irrespective of conformational state (Figure 5b, Supplementary Figures 9–12). These results confirm that distal mutations alter the LEF within the active site.

**Figure 5.**
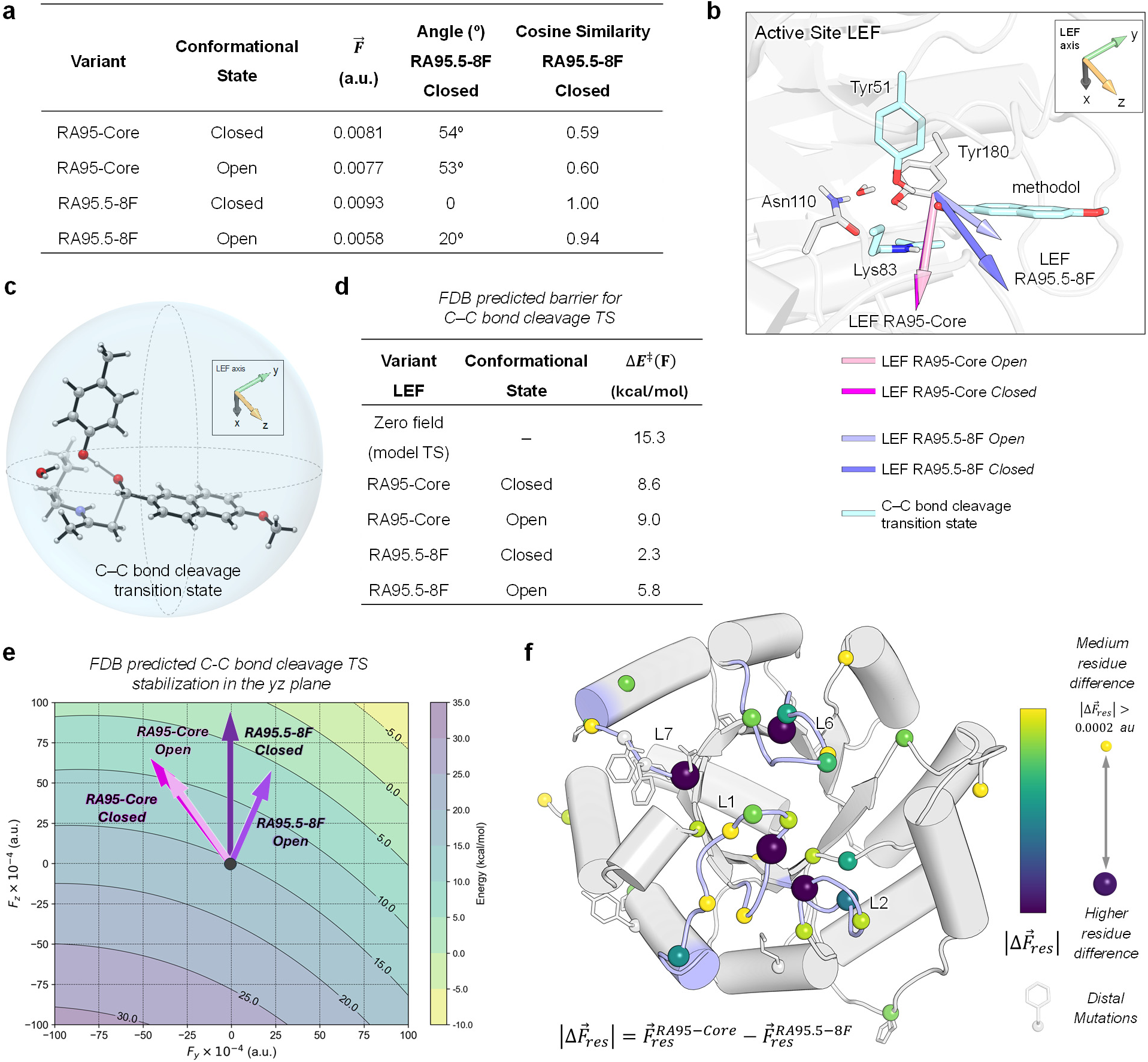
Distal mutations alter the local electric field (LEF) to stabilize the C–C bond cleavage transition state (TS). (a) Calculated magnitude and orientation of the active-site LEF 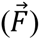 for retro-aldolase variants in various conformational states. LEF orientation is described relative to the Closed state of RA95.5-8F, using the angle and cosine similarity measure (inner product space) between LEF vectors. (b) Active-site structure showing LEF vectors for each conformational state and variant. The theozyme TS model, including Lys83, Tyr51, and the methodol substrate, is shown in cyan sticks. The rest of the catalytic tetrad is depicted as gray sticks. The aligned theozyme structure corresponds to RA95-Core Closed (see Methods). (c) Optimized theozyme TS model for the C–C bond cleavage step. (d) Ideal energy barrier (ΔE^‡^) for the rate-limiting C–C bond cleavage step, based on the theozyme model, calculated under zero-field conditions and with LEFs corresponding to those determined for each enzyme variant and conformational state (panel a). (e) Two-dimensional representation of the chemical barriers for the C–C bond cleavage step estimated from the theozyme model in terms of LEFs along the *y* and *z*-axes using the FDB approach. This analysis shows that LEFs generated in the RA95.5-8F active site have an optimal orientation for TS stabilization. (f) Residues contributing the most to LEF changes (from RA95-Core to RA95.5-8F) are shown as coloured spheres. Sphere size and colour indicate the magnitude of the contribution. The most significant changes in LEF arise from residues located on flexible loops (L1, L2, L6, L7), rather than directly from the distal mutation sites (white spheres). The protein scaffold corresponds to RA95-Core Closed.

To assess the functional implications of these differences, we applied the field-dependent energy barrier method (*34*) to a truncated transition state model of the C–C bond cleavage step (Figure 5c). This analysis revealed that the LEF generated in RA95.5-8F intrinsically reduces the energy barrier for the C–C bond cleavage step by 3–6 kcal mol^−1^ compared to the LEF in RA95-Core (Figure 5d,e), showcasing the importance of the orientation of the field in stabilizing that particular transition state. Residue-level analysis further showed that the largest changes in LEF contributions arose from residues on flexible loops, rather than directly from the distal mutation sites (Figure 5f, Supplementary Figure 13). These findings indicate that distal mutations enhance catalysis by modulating enzyme conformational dynamics, which reorient the LEF to optimize electrostatic preorganization of the active site. This mechanism is consistent with the observed increase in catalytic efficiency for C–C bond cleavage in RA95.5-8F, highlighting the critical role of distal mutations in shaping electrostatic preorganization through their influence on enzyme dynamics.

## Discussion

Understanding how distal amino-acid residues influence catalytic function is critical for advancing our knowledge of enzyme catalysis (*35*). In this study, we investigated the effects of distal mutations with and without the accompanying active-site mutations that were co-selected for enhanced catalysis during a directed evolution campaign. Our results indicate that in the context of both the optimized active site (found in RA95-Core and RA95.5-8F) and the original, suboptimal active site (found in RA95 and RA95-Shell), distal mutations altered loop structure and dynamics, facilitating opening of the active site. In the presence of the optimized active site, this enhanced opening increased the rate of product release by 4-fold, contributing to the 14-fold higher *k*_cat_ of RA95.5-8F. However, in the context of the suboptimal RA95 active site, increased opening did not enhance *k*_cat_, likely because its C–C bond cleavage rate (*k* = 0.00011 s^−1^) (*26*) is three orders of magnitude slower than RA95-Core (*k* = 0.43 s^−1^). Even if the rate of product release in RA95-Shell matched that of RA95.5-8F, C–C bond cleavage would remain rate-limiting, preventing significant improvement in *k*_cat_. Interestingly, the penultimate variant in the RA95 evolutionary trajectory, RA95.5-8, exhibits a *k*_cat_ similar to RA95-Core (*k*_cat_ = 0.36 s^−1^) but demonstrates a 6-fold slower product release (*k* = 0.21 s^−1^) (*26*). RA95.5-8 contains four of the ten distal mutations identified during evolution (Supplementary Table 1), which are absent in RA95-Core, suggesting that the remaining six mutations primarily drive the accelerated product release. Together, these data emphasize that while specific distal mutations can alter the conformational ensemble in similar ways when introduced on different active sites, their effects on catalytic function are dependent on epistatic interactions with active-site mutations (*36*).

Distal mutations also accelerated C–C bond cleavage by two orders of magnitude when introduced into RA95-Core, shifting the rate-limiting step from C–C bond cleavage to product release. The faster chemical transformation observed in RA95.5-8F, despite it having the same active-site residues as RA95-Core, indicates that subtle structural and dynamic changes caused by distal mutations further optimize the active-site environment for efficient catalysis. Previous molecular dynamics studies of RA95 variants have demonstrated that distal mutations, in conjunction with active-site mutations, stabilize catalytically competent conformations, shifting populations toward productive sub-states in both the Schiff base intermediate and unbound enzymes (*6*). While this effect was attributed to conformational changes observed in active-site loops, our results demonstrate that the shift toward catalytically competent conformations during evolution can be ascribed to distal mutations. In addition, our data indicates that these mutations alter the local electric field in the active site through dynamic changes, further enhancing catalysis in RA95.5-8F compared to RA95-Core, despite their identical active-site sequences. Overall, our findings underscore the multifaceted role of distal mutations in enhancing enzyme efficiency by modulating loop dynamics and optimizing the active-site environment, geometrically and electrostatically, to support an efficient catalytic cycle.

These results challenge traditional enzyme design strategies that focus on optimizing active-site interactions for transition-state stabilization (*23–25*). Our work demonstrates that, even if RA95-Core’s active site were perfectly designed, it would not be the most active enzyme producible on that scaffold, because distal mutations increase the rate of both C–C bond cleavage and product release. Recently, deep learning methods were employed to design *de novo* retro-aldolases by constructing an entire protein scaffold around the RA95.5-8F catalytic tetrad (*37*), yielding the most active computationally designed retro-aldolases to date. However, the best variant from this study (*k*_cat_ = 0.031 s^−1^) is still 150-fold less active than RA95.5-8F, despite having a comparable *K*_M_ (100 µM). Furthermore, crystal structures revealed that accurate positioning of the catalytic tetrad alone does not ensure high catalytic efficiency; indeed, some of the most structurally accurate designs were among the least active (*37*). Therefore, integrating insights from our findings into deep learning frameworks could enhance enzyme design by prioritizing allosteric effects that tune dynamic flexibility and optimize active-site electric fields while also improving structural accuracy of the active site.

More broadly, our results provide important insights into both direct and epistatic effects of distal mutations on the enzyme catalytic cycle. When product release is the rate-limiting step, as is the case for many natural enzymes (*38–40*), optimization of large-scale structural and dynamic changes across the entire protein scaffold becomes necessary to achieve further rate enhancement. Collectively, our findings shed new light on how distal regions allosterically influence the catalytic cycle to drive catalytic efficiency, offering insights that could guide the design of more efficient *de novo* enzymes (*41*), improve our understanding of how disease mutations disrupt enzyme function (*42, 43*), and elucidate the physical underpinnings of epistasis in shaping the evolutionary trajectories of natural enzymes (*36*).

## Materials and Methods

### Protein expression and purification

Codon-optimized (*E. coli*) and His-tagged (C-terminus) retro-aldolase genes (Supplementary Tables 3–4) cloned into the pET-29b(+) vector via *Nde*I and *Xho*I restriction sites were obtained from Twist Bioscience. Enzymes were expressed in *E. coli* BL21-Gold (DE3) cells (Agilent) using lysogeny broth (LB) supplemented with 50 µg mL^−1^ kanamycin. Cultures were grown at 37 °C with shaking (220 rpm) to an optical density of approximately 0.6 at 600 nm. Protein expression was then induced with 1 mM isopropyl β-D-1-thiogalactopyranoside and cells were incubated for 16 hours at 18 °C with shaking (220 rpm). Cells were harvested by centrifugation, resuspended in 10 mL lysis buffer (5 mM imidazole in 100 mM potassium phosphate buffer, pH 8.0), and lysed with an EmulsiFlex-B15 cell disruptor (Avestin). Proteins were purified by immobilized metal affinity chromatography using Ni–NTA agarose (Qiagen) pre-equilibrated with lysis buffer in individual Econo-Pac gravity-flow columns (Bio-Rad). Columns were washed twice with lysis buffer. Bound proteins were eluted with 250 mM imidazole in 100 mM potassium phosphate buffer (pH 8.0) and exchanged into 25 mM HEPES buffer (pH 7.5) supplemented with 100 mM NaCl using Econo-Pac 10DG desalting pre-packed gravity-flow columns (Bio-Rad). Protein samples for crystallography were further subjected to purification by gel filtration in 20 mM potassium phosphate buffer (pH 7.4) and 50 mM NaCl using an ENrich SEC 650 size-exclusion chromatography column (Bio-Rad). Purified protein samples were quantified by measuring the absorbance at 280 nm and applying Beer-Lambert’s law using calculated extinction coefficients obtained from the ExPAsy ProtParam tool (https://web.expasy.org/protparam/).

### Steady-state kinetics

Steady-state kinetic assays were carried out at 29 °C in 25 mM HEPES (pH 7.5) supplemented with 100 mM NaCl. Triplicate 200-µL reactions with varying concentrations of freshly-prepared racemic 4-hydroxy-4-(6-methoxy-2-naphthyl)-2-butanone (methodol) (Achemica) dissolved in acetonitrile (2.7% final concentration) were initiated by the addition of 180 µM RA95, 0.1 µM RA95.5-8F, 2 µM RA95-Core, 120 µM RA95-Shell, 20 µM RA95-Core-Y51F, 4 µM RA95-Core-N110S or 4 µM RA95-Core-Y180F. Product (6-methoxy-2-napthaldehyde) formation was monitored spectrophotometrically at 350 nm (ε = 5,970 M^−1^ cm^−1^) (*14*) in individual wells of 96-well plates (Greiner Bio-One) using a SpectraMax 384Plus plate reader (Molecular Devices). Path lengths for each well were calculated ratiometrically using the difference in absorbance of 25 mM HEPES (pH 7.5) supplemented with 100 mM NaCl and 2.7% acetonitrile at 900 and 975 nm (29 ° C). Linear phases of the kinetic traces were used to measure initial reaction rates. *k*_cat_ and *K*_M_ were determined by fitting the data to the Michaelis-Menten model v_0_ = *k*_cat_[E_0_][S]/(*K*_M_+[S]) in GraphPad Prism 5.

### Kinetic solvent viscosity effects

The effects of solvent viscosity on steady-state kinetic parameters (*27*) *k*_cat_/*K*_M_ and *k*_cat_ were determined at 29 °C in 25 mM HEPES buffer (pH 7.5) supplemented with 100 mM NaCl using sucrose as viscogen at different concentrations (0, 20, 28, 33 % weight/volume). Corresponding viscosities (η) were approximated from published viscosity data of sucrose solutions (*44*). Steady-state kinetic assays were performed as described above using 320–800 nM and 2–4 µM of purified RA95.5-8F and RA95-Core, respectively. Initial rates were determined and fitted to the Michaelis-Menten equation to calculate *k*_cat_/*K*_M_ and *k*_cat_ values. The reference value at 0 % sucrose was divided by those obtained at different η and plotted against the relative buffer viscosity η_rel_ to give the corresponding slopes (Figure 4a,d). Rate constants for the chemical transformation (*k*_3_) and product dissociation (*k*_4_) (Scheme 1) were calculated using *k*_cat_ and the slope (m) according to equations 1 and 2:

**Scheme 1.**
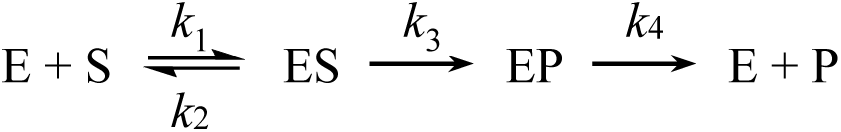
Kinetic mechanism for an irreversible enzymatic (E) reaction with a single substrate (S) and product (P) showing rate constants for substrate association (*k*_1_) and dissociation (*k*_2_), the chemical transformation (*k*_3_), and product dissociation (*k*_5_).

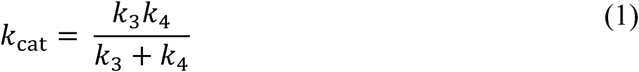

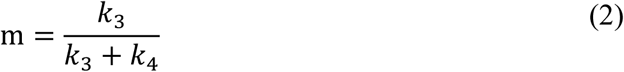

### pH rate profile

Steady-state kinetic assays for pH rate profile determination were carried out at 29 °C in Britton-Robinson buffer (40 mM boric acid, 40 mM phosphoric acid, 40 mM acetic acid) at varying pH values. Triplicate 200-µL reactions with varying concentrations of freshly prepared racemic methodol (Achemica) dissolved in acetonitrile (2.7% final concentration) were initiated by the addition of the enzyme. Product (6-methoxy-2-napthaldehyde) formation was monitored spectrophotometrically at 350 nm (ε = 5,970 M^−1^ cm^−1^) (*14*) in individual wells of 96-well plates (Greiner Bio-One) using a SpectraMax 384Plus plate reader (Molecular Devices). Path lengths for each well were calculated ratiometrically using the difference in absorbance of the Britton-Robinson buffer supplemented with 2.7 % acetonitrile at 900 and 975 nm (29 °C). Linear phases of the kinetic traces were used to measure initial reaction rates. Initial reaction rates were fitted to the linear portion of the Michaelis-Menten model v_0_ = (*k*_cat_/*K*_M_) [E_0_], and *k*_cat_/*K*_M_ was deduced from the slope. To determine p*K*a values, *k*_cat_/*K*_M_ data were fitted in GraphPad Prism 5 to the following equation using nonlinear least squares regression: (*k*_cat_/*K*_M_)_obs_ = (*k*_cat_/*K*_M_)_max_/(1+10^p*K*a1–pH^ + 10^p*K*a2–pH^).

### Circular dichroism (CD) and thermal denaturation assays

CD measurements were performed with a Jasco J-815 spectrometer using 300-µL aliquots of each retro-aldolase sample at a concentration of 5 µM in 10 mM sodium phosphate buffer (pH 7.0) in a 1-mm path-length quartz cuvette (Jasco). For structural characterization of protein folds, CD spectra were acquired from 200 to 250 nm at 20 °C, sampled every 1 nm at a rate of 10 nm min^−1^. Three scans were acquired and averaged for each sample. For thermal denaturation assays, samples were heated at a rate of 1 °C per minute, and ellipticity at 222 nm was measured every 0.2 °C. Melting temperatures were determined by fitting the data to a two-term sigmoid function with correction for pre- and post-transition linear changes in ellipticity as a function of temperature (*45*). Data were fitted to equations 3 through 6 using nonlinear least-squares regression in GraphPad Prism 5, where θ_F_ is the ellipticity when 100% folded, θ_U_ is the ellipticity when 100% unfolded, c_F_ is the linear correction for pre-transition changes in ellipticity, c_U_ is the linear correction for post-transition changes in ellipticity, ΔH_U_ is the enthalpy of unfolding, k is the folding constant, F is the fraction folded, and θ is the ellipticity at temperature T.

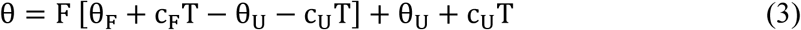

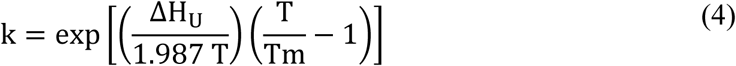

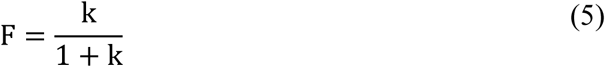

### Crystallization

Purified protein samples were concentrated to 10–15 mg mL^−1^ using Amicon Ultracel-3K centrifugal filter units (EMD Millipore). Crystals were obtained by the hanging-drop vapour diffusion method at 293 K in drops prepared by mixing 1 μL of protein solution with 1 μL of the mother liquor and sealing the drop inside a reservoir containing an additional 500 μL of the mother liquor solution. The mother liquor solution that yielded the crystals of RA95 used for X-ray data collection contained 0.1 M sodium acetate (pH 5.2) and 3.1 M NaCl with a protein solution concentration of 7 mg mL^−1^. The mother liquor solution that yielded crystals of RA95-Shell used for X-ray data collection contained 0.1 M sodium acetate (pH 4.4) and 19 % weight/volume PEG 3,000 with a protein solution concentration of 6 mg mL^−1^.

### X-ray data collection and processing

Crystals were mounted on polyimide loops and sealed using a MicroRT tubing kit (MiTeGen). Single-crystal X-ray diffraction data were collected on beamline 8.3.1 at the Advanced Light Source. The beamline was equipped with a Pilatus3 S 6M detector (Dectris) and was operated at a photon energy of 11111 eV. Crystals were maintained at 277 and 280 K for RA95 and RA95-Shell, respectively, throughout the course of data collection. Multiple data sets were collected for each variant either from different crystals or from different regions of larger crystals. X-ray data were processed using the Xia2 software (*46*), which performed indexing, integration, and scaling with XDS and XSCALE (*47*), followed by merging with Pointless (*48*).

### Structure determination

Initial phase information for calculation of electron density maps was obtained by molecular replacement using the program Phaser (*49*), as implemented in v1.13.2998 of the PHENIX suite (*50*). The previously published RA95 structure with bound inhibitor (PDB ID: 4A29) (*14*) was used as the molecular replacement search model. All members of the RA95-series of enzymes crystallized in the same crystal form, containing one copy of the molecule in the crystallographic asymmetric unit. Next, we performed iterative steps of manual model rebuilding followed by refinement of atomic positions, atomic displacement parameters, and occupancies using a translation-libration-screw (TLS) model, a riding hydrogen model, and automatic weight optimization. All model building was performed using Coot 0.8.9.236 (*51*) and refinement steps were performed with *phenix.refine* (*52*) within the PHENIX (v1.13-2998) suite. Further information regarding model building and refinement are presented in Supplementary Table 2.

### Unconstrained molecular dynamics (MD)

Microsecond timescale MD simulations were performed in triplicate using the Amber 2020 software (http://ambermd.org/) with the AMBER19SB force field (*53*). Long-range electrostatics (>10 Å) were modeled using the particle mesh Ewald method (*54*), and a time step of 2 fs was used for the production phase. Unbound crystal structures of RA95 (PDB ID: 9MYA), RA95-Shell (PDB ID: 9MYB), and RA95.5-8F (PDB ID: 5AOU) (*15*) were used for molecular dynamics. Missing residues (58–63) of the RA95.5-8F crystal structure were modelled using MODELLER 10.4 (*55*) by selecting only the missing residues using the *AutoModel* class. The unbound structure of RA95-Core was generated from the unbound crystal structure of RA95 by introducing mutations with the *sequenceDesign.py* app in the Triad protein design software v2.1.2 (Protabit LLC, Pasadena, CA) (*56*), which optimized rotameric configurations of the active site. Amino acid protonation states were predicted using the H++ server (http://biophysics.cs.vt.edu/H++) at pH 7.0. Prior to molecular dynamics, the structures were parameterized using the LEaP program from the AMBER suite. The protein molecule was placed in a dodecahedral box with periodic boundary conditions where the distance between the protein surface and the box edges were set to 10 Å. After the addition of explicit TIP3P water molecules (*57*), charges on protein atoms were neutralized with Na^+^ and Cl^−^ counter-ions at a concentration of 0.15 M. The structures were then energy minimized with the steepest descent method to a target maximum force of 1000 kJ mol^−1^ nm^−1^. Before equilibration, the system was heated to a target temperature of 300 K for 240 ps. The system was then equilibrated under an NPT ensemble for 10 ns with constant pressure and temperature of 1 bar and 300 K, respectively, using the Berendsen barostat (*58*). A second equilibration step under an NVT ensemble was performed for 10 ns at a temperature of 300 K using the Langevin temperature equilibration scheme. 1000-ns production runs were initiated from the final snapshot of the NVT equilibration. Principal component analysis and k-means clustering were done with the *pyEMMA* software (*59*). Snapshots separated by 20 ps along the production trajectories were extracted for principal component analysis. After partitioning of the trajectories by k-means clustering, 1,500 snapshots separated by 2 ns were used for the mean loop distances and the histograms in Figure 3.

### Electric field calculations

Centroid structures corresponding to the open and closed states obtained from clusterization of MD trajectories were aligned with the crystal structure of RA95.5-8F with a bound covalent inhibitor (PDB: 5AN7). An arbitrary point to describe the electric field in the active site was defined to coincide with the position occupied by the hydroxyl oxygen atom of the inhibitor molecule in the 5AN7 structure. The CPPTRAJ module from AmberTools was used to strip water and ions from the selected snapshots of the centroid structures. The strength and direction of the local electric field at the selected point was calculated considering the classic definition of the electrostatic forces between particles in a system using Coulomb’s Law. In this context, the electric field 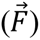 exerted by N atoms at a given point in the simulation box can be estimated as defined by equation 6:

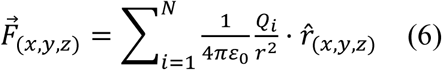

in which *ε_0_* is the permittivity of vacuum, *Q_i_* is the partial charge of atom *i*, *r* is the distance between atom *i* and the (*x,y,z*) point in space and *r̂* is the unit vector of the distance. The TUPÃ software (*60*) and the same Amber derived charges used for the MD simulations were utilized. The entire protein was considered except for Lys83 and Tyr51 catalytic residues, which are directly participating in the chemical step and are already considered in the theozyme truncated model (see below). The pyTUPAmol plugin for PyMOL was used to plot the electric field vector of the centroid and cluster ensembles.

The representative nature of the selected arbitrary point was proven by analyzing a grid of points in the active sites of the studied systems, confirming that these points effectively describe the trend of the LEF generated at each active site cavity (Supplementary Figures 10–11). A cubic box was created centered at the hydroxyl oxygen, extending 2 Å in each direction along the X, Y, and Z axes, with a grid spacing of 1 Å. This resulted in a total of 125 points. The electric field was calculated using the TUPÃ software at each grid point in the active site (Supplementary Figure 12). These analyses support that the selected point correctly describes the behavior of the local electric field in this region of the active site pocket.

### Quantum mechanics theozyme calculations

The crystal structure of RA95.5-8F with bound covalent inhibitor (PDB: 5AN7) was used as a model system for preparing the theozyme truncated model. The inhibitor along with the Lys83 and Tyr51 side chains were extracted, and modifications were manually made to create a theozyme truncated model of the carbinolamine intermediate and to model the C–C bond cleavage step. Quantum mechanical (QM) density functional theory (DFT) calculations were preformed using Gaussian16 (*61*). The unrestricted hybrid (U)B3LYP functional (*62–64*) was used with an ultrafine integration grid, and including the CPCM polarizable conductor model (dichloromethane, ε = 8.93) (*65, 66*) to have an estimation of the dielectric permittivity in the enzyme active site (*67*). 6-31G(d) basis set was used for all atoms. All optimized stationary points were characterized as minima using frequency calculations, including transition states which show a single imaginary frequency that describes the corresponding reaction coordinate. IRC calculations were performed to ensure that optimized transition states connect the expected reactants and products. Figures were rendered using CYLview (http://www.cylview.org). Dipole moments, and (hyper)polarizabilities were obtained at the same level using the "*polar*" keyword in Gaussian16.

Field dependent energy barriers (FDB) (*34*) were calculated using the strategy proposed by Torrent-Sucarrat, Luis and co-workers, with the truncation of the Taylor expansion around free-field energy at the third-order correction (given by the first hyperpolarizability, β, see eq. 7), and the open-access script provided by the authors (https://github.com/pau-besalu/FDB).

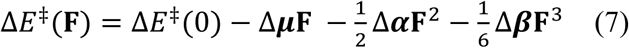

To validate the predictions from the FDB, we have also computed the effect of explicit external static electric fields on the energy barriers using Gaussian16. Optimized *theozyme* transition state and reactant structures were manually aligned with the inhibitor bound in RA95.5-8F and catalytic Lys83 and Tyr51 residues, which was already aligned also with the the RA95.5-8F, RA95-Shell and RA95-Core centroid structures. The strength and direction of the electric fields 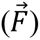 estimated at the active site of each of these structures (see electric field methodology section) were considered, obtaining a perfect agreement between FDB predicted energy barriers and those obtained from explicit electric field calculations.

## Data availability

Structure coordinates for all retro-aldolases have been deposited in the RCSB Protein Data Bank with the following accession codes: RA95 (PDB ID: 9MYA) and RA95-Shell (PDB ID: 9MYB). Source data are provided with this paper. Other relevant data are available from the corresponding authors upon reasonable request.

## Acknowledgements

R.A.C. acknowledges grants from the Natural Sciences and Engineering Research Council of Canada (RGPIN-2021-03484 and RGPAS-2021-00017) and the Canada Foundation for Innovation (26503). R.A.C. and M.C.T. acknowledge a joint grant from the Human Frontier Science Program (RGP0004/2022). S.E.H. is the recipient of an Ontario Graduate Scholarship. M.G.-B. and F.F. acknowledge the Spanish Ministry of Science and Innovation MICINN for PID2022-141676NB-I00 and TED2021-130173B-C42 projects, and RYC2020-028628-I (M.G.-B.) and RYC2020-029552-I (F.F.) grants, and the Generalitat de Catalunya for 2021SGR00623 and 2021SGR00487 projects. A.E.J. is supported by FPI2023_PRE49-2023 predoctoral fellowship. This research was enabled in part by support provided by Compute Ontario (www.computeontario.ca) and the Digital Research Alliance of Canada (alliancecan.ca). Beamline 8.3.1 at the Advanced Light Source is operated by the University of California San Francisco with generous support from the National Institutes of Health (R01 GM124149 for technology development and P30 GM124169 for user support), and the Integrated Diffraction Analysis Technologies program of the US Department of Energy Office of Biological and Environmental Research. The Advanced Light Source (Berkeley, CA) is a national user facility operated by Lawrence Berkeley National Laboratory on behalf of the US Department of Energy under contract number DE-AC02-05CH11231, Office of Basic Energy Sciences. The contents of this publication are solely the responsibility of the authors and do not necessarily represent the official views of NIGMS or NIH.

## Author Contributions

R.A.C. and S.E.H. designed the project. S.E.H. and A.M. performed protein crystallization. M.C.T. performed X-ray data collection and processing. R.A.C. performed refinement of crystal structures. N.Z. designed the RA95-Core and RA95-Shell variants. C.K. performed kinetic solvent viscosity effect experiments. M.G.-B. and F.F. designed the electric field analysis strategy and analyzed the data, and A.E.J. performed the calculations. S.E.H. completed all other experimental and computational experiments. R.A.C. and S.E.H. wrote the manuscript with input from the other authors. All authors revised the manuscript.

