## Supplementary Information for "Distal mutations in a designed retro-aldolase alter loop dynamics to shift and accelerate the rate-limiting step"

**This file includes:**

Supplementary Tables 1–4

Supplementary Figures 1–13

### Supplementary Table 1. Mutations in RA95 variants

| Enzyme | # of mutations from RA95 | Mutations from RA95 |
|---|---|---|
| RA95 | - | - |
| RA95.5 | 6 | V51Y, E53S, T83K, M180F, R182M, D183N |
| RA95.5-5 [a] | 11 | R23H, R43S, V51Y, E53T*, T83K, T95M, S110N, G178S, M180F, R182M, D183N |
| RA95.5-8 [a] | 13 | R23H, V51Y, E53T*, F72Y, T83K, T95M, S110N, K135N, G178V*, M180F, R182M, D183N, G212D |
| RA95.5-8F [a] | 22 | R23H, V51Y, E53L**, F72Y, R75P, T83K, N90D, T95M, S110N, K135E*, S151G, G178T**, M180Y*, R182M, D183N, A209P, K210L, G212D, I213F, S214F, R216P, L231M |
| RA95-Core | 12 | V51Y, E53L, T83K, N90D, S110N, K135E, G178T, M180Y, R182M, D183N, K210L, L231M |
| RA95-Shell | 10 | R23H, F72Y, R75P, T95M, S151G, A209P, G212D, I213F, S214F, R216P |
| RA95-Core-Y51F | 12 | V51F, E53L, T83K, N90D, S110N, K135E, G178T, M180Y, R182M, D183N, K210L, L231M |
| RA95-Core-N110S | 11 | V51Y, E53L, T83K, N90D, K135E, G178T, M180Y, R182M, D183N, K210L, L231M |
| RA95-Core-Y180F | 12 | V51Y, E53L, T83K, N90D, S110N, K135E, G178T, M180F, R182M, D183N, K210L, L231M |

[a] One and two asterisks indicate positions that were mutated two or three times, respectively, during the RA95 evolutionary trajectory.

**Supplementary Table 2. Crystallography data and refinement statistics**

| | **RA95** | **RA95-Shell** |
|---|---|---|
| PDB ID | 9MYA | 9MYB |
| Crystallizations conditions | 0.1 M sodium acetate pH 5.2<br>3.1 M NaCl<br>7 mg $mL^{-1}$ protein | 0.1 M sodium acetate pH 4.4<br>19% PEG 3000<br>6 mg $mL^{-1}$ protein |
| Protein buffer | 20 mM potassium phosphate pH 7.4<br>50 mM NaCl | 20 mM potassium phosphate pH 7.4<br>50 mM NaCl |
| **Data collection** [a] | | |
| Temperature (K) | 277 | 280 |
| Resolution (Å) | 49.00–1.89 | 51.66–1.77 |
| Space group | P 21 21 2 | P 21 21 2 |
| *Cell params.* | | |
| a b c (Å) | 97.995<br>65.156<br>44.377 | 85.638<br>64.804<br>41.034 |
| α β γ (°) | 90<br>90<br>90 | 90<br>90<br>90 |
| Chains per asymm. unit | 1 | 1 |
| $R_{pim}$ | 0.093 (0.464) | 0.098 (0.648) |
| $CC_{1/2}$ | 0.988 (0.576) | 0.991 (0.391) |
| I/σI | 4.9 (0.9) | 7.0 (1.0) |
| Completeness (%) | 98.6 (97.3) | 100.0 (100.0) |
| Multiplicity | 6.4 (6.7) | 6.3 (5.9) |
| Wilson B-factor ($Å^2$) | 18.820 | 17.570 |
| # unique reflections | 23151<br>(1137) | 22968<br>(1127) |
| **Refinement** | | |
| R work/free | 0.1588/0.1969 | 0.1869/0.2184 |
| *No. atoms* | | |
| Protein | 2109 | 2073 |
| Ligand | 1 | 0 |
| Water | 109 | 77 |
| *Averaged B-factors ($Å^2$)* | | |
| Protein | 30.53 | 30.46 |
| Ligands | 29.68 | – |
| Water | 34.19 | 32.22 |
| *RMSD* | | |
| bond lengths (Å) | 0.011 | 0.003 |
| bond angles (°) | 1.007 | 0.525 |
| *Molprobity statistics* | | |
| Ramachand. outliers (%) | 0.00 | 0.00 |
| Ramachand. allowed (%) | 1.63 | 0.82 |
| Ramachan. favored (%) | 98.37 | 99.18 |
| Rotamer outliers (%) | 0.00 | 0.00 |
| MolProbity clashscore | 2.09 | 0.95 |

[a] Highest resolution shell is shown in parentheses.

### Supplementary Table 3. Amino-acid sequences of RA95 variants

| Enzyme | Sequence [a] |
|---|---|
| RA95 | PRYLKGWLEDVVQLSLRRPSVRASRQRPIISLNERILEFNKRNITAIIAVYERKSPSGLDVERDPIEYAKFMERYAV<br>GLSITTEEKYFNGSYETLRKIASSVSIPILMSDFIVKESQIDDAYNLGADTVLLIVKILTERELESLLEYARSYGME<br>PLILINDENDLDIALRIGARFIGIMSRDFETGEINKENQRKLISMIPSNVVKVAKLGISERNEIEELRKLGVNAFLI<br>SSSLMRNPEKIKELIEGSLEHHHHHH |
| RA95.5-8F | PRYLKGWLEDVVQLSLRRPSV**H**ASRQRPIISLNERILEFNKRNITAIIA**Y**Y**L**RKSPSGLDVERDPIEYAK**Y**ME**P**YAV<br>GLSI**K**TEEKYF**D**GSYE**M**LRKIASSVSIPILM**N**DFIVKESQIDDAYNLGADTVLLIV**E**ILTERELESLLEYAR**G**YGME<br>PLILINDENDLDIALRIGARFI**T**I**Y**S**MN**FETGEINKENQRKLISMIPSNVVKV**PL**L**DFF**E**P**NEIEELRKLGVNAF**M**I<br>SSSLMRNPEKIKELIEGSLEHHHHHH |
| RA95-Shell | PRYLKGWLEDVVQLSLRRPSV**H**ASRQRPIISLNERILEFNKRNITAIIAVYERKSPSGLDVERDPIEYAK**Y**ME**P**YAV<br>GLSITTEEKYFNGSYE**M**LRKIASSVSIPILMSDFIVKESQIDDAYNLGADTVLLIVKILTERELESLLEYAR**G**YGME<br>PLILINDENDLDIALRIGARFIGIMSRDFETGEINKENQRKLISMIPSNVVKV**P**KL**DFF**E**P**NEIEELRKLGVNAFLI<br>SSSLMRNPEKIKELIEGSLEHHHHHH |
| RA95-Core | PRYLKGWLEDVVQLSLRRPSVRASRQRPIISLNERILEFNKRNITAIIA**Y**Y**L**RKSPSGLDVERDPIEYAKFMERYAV<br>GLSI**K**TEEKYF**D**GSYETLRKIASSVSIPILM**N**DFIVKESQIDDAYNLGADTVLLIV**E**ILTERELESLLEYARSYGME<br>PLILINDENDLDIALRIGARFI**T**I**Y**S**MN**FETGEINKENQRKLISMIPSNVVKVA**L**LGISERNEIEELRKLGVNAF**M**I<br>SSSLMRNPEKIKELIEGSLEHHHHHH |
| RA95-Core-Y51F | PRYLKGWLEDVVQLSLRRPSVRASRQRPIISLNERILEFNKRNITAIIA**F**Y**L**RKSPSGLDVERDPIEYAKFMERYAV<br>GLSI**K**TEEKYF**D**GSYETLRKIASSVSIPILM**N**DFIVKESQIDDAYNLGADTVLLIV**E**ILTERELESLLEYARSYGME<br>PLILINDENDLDIALRIGARFI**T**I**Y**S**MN**FETGEINKENQRKLISMIPSNVVKVA**L**LGISERNEIEELRKLGVNAF**M**I<br>SSSLMRNPEKIKELIEGSLEHHHHHH |
| RA95-Core-N110S | PRYLKGWLEDVVQLSLRRPSVRASRQRPIISLNERILEFNKRNITAIIA**Y**Y**L**RKSPSGLDVERDPIEYAKFMERYAV<br>GLSI**K**TEEKYF**D**GSYETLRKIASSVSIPILM**N**DFIVKESQIDDAYNLGADTVLLIV**E**ILTERELESLLEYARSYGME<br>PLILINDENDLDIALRIGARFI**T**I**Y**S**MN**FETGEINKENQRKLISMIPSNVVKVA**L**LGISERNEIEELRKLGVNAF**M**I<br>SSSLMRNPEKIKELIEGSLEHHHHHH |
| RA95-Core-Y180F | PRYLKGWLEDVVQLSLRRPSVRASRQRPIISLNERILEFNKRNITAIIA**Y**Y**L**RKSPSGLDVERDPIEYAKFMERYAV<br>GLSI**K**TEEKYF**D**GSYETLRKIASSVSIPILMSDFIVKESQIDDAYNLGADTVLLIV**E**ILTERELESLLEYARSYGME<br>PLILINDENDLDIALRIGARFI**T**I**F**S**MN**FETGEINKENQRKLISMIPSNVVKVA**L**LGISERNEIEELRKLGVNAF**M**I<br>SSSLMRNPEKIKELIEGSLEHHHHHH |

[a] Mutations from RA95 are highlighted in bold and underlined. All sequences contain a 6×His-tag at the C-terminus.

**Supplementary Table 4. DNA sequences of RA95 variants.**

| Enzyme | Sequence |
|---|---|
| RA95 | CCGCGTTACTTGAAAGGATGGCTTGAAGATGTGGTTCAATTGTCGTTACGCCGCCCATCGGTCCGTGCCAGTCGTCAGCGTCC<br>CATTATCTCCCTGAACGAGCGTATCTTGGAGTTTAACAAGCGTAATATTACGGCTATCATCGCCGTGTATGAGCGTAAGTCGC<br>CTTCCGGTCTGGACGTTGAACGCGATCCAATTGAGTACGCCAAATTTATGGAGCGTTATGCGGTGGGTTTGTCGATTACGACT<br>GAAGAGAAGTATTTCAACGGCTCATACGAAACTTTGCGTAAGATTGCGTCGTCGGTCAGCATCCCCATCCTGATGTCGGATTT<br>CATCGTAAAAGAGAGCCAGATCGACGATGCATACAATCTGGGTGCTGACACAGTTCTTCTGATTGTGAAGATCTTAACAGAAC<br>GTGAGTTAGAGTCCTTGCTTGAATACGCGCGTAGCTACGGCATGGAACCTTTGATTCTTATCAACGACGAAAATGATCTTGAT<br>ATCGCGTTACGTATTGGTGCGCGTTTCATCGGGATTATGTCGCGCGATTTCGAGACCGGTGAGATCAACAAGGAAAATCAACG<br>CAAGCTTATTAGCATGATCCCTTCCAATGTTGTGAAGGTTGCAAAATTGGGCATTTCGGAGCGCAACGAGATCGAAGAGCTGC<br>GTAAATTGGGAGTCAATGCATTCTTGATCTCCAGCTCTCTGATGCGCAATCCTGAGAAAATCAAGGAGTTAATTGAAGGTAGC<br>CTGGAGCACCACCATCACCACCATTAA |
| RA95.5-8F | CCTCGCTATTTGAAGGGGTGGCTGGAAGACGTAGTACAACTCTCATTGCGTCGCCCATCAGTTCATGCGAGTCGTCAACGCCC<br>GATTATCTCATTGAACGAACGCATTCTTGAGTTCAATAAGCGTAATATCACCGCCATCATCGCGTATTACCTTCGCAAGAGTC<br>CTAGTGGTCTTGACGTAGAACGCGATCCGATTGAGTACGCCAAGTACATGGAACCGTACGCGGTAGGTTTAAGCATTAAGACC<br>GAAGAGAAGTATTTTGACGGCTCCTACGAAATGTTACGCAAAATCGCCTCTAGTGTTAGTATTCCAATCCTCATGAACGATTT<br>CATCGTTAAGGAATCACAGATCGACGATGCCTACAATTTAGGTGCAGACACCGTGTTGCTTATTGTCGAAATTTTAACCGAAC<br>GCGAGTTAGAGTCTCTCCTTGAATACGCACGTGGCTACGGTATGGAGCCCTTAATTCTGATTAATGATGAGAATGATCTCGAT<br>ATCGCGCTGCGCATCGGCGCTCGCTTCATCACAATCTATTCCATGAATTTTGAGACGGGTGAAATTAATAAAGAGAATCAACG<br>CAAGTTAATTAGCATGATTCCGAGCAACGTGGTGAAGGTGCCCCTTCTGGACTTCTTCGAGCCCAACGAGATTGAAGAGTTAC<br>GTAAGCTCGGCGTGAATGCGTTCATGATTTCCTCTAGCCTTATGCGTAATCCCGAGAAAATTAAAGAGTTAATCGAGGGGTCC<br>TTAGAGCATCATCACCACCACCACTGA |
| RA95-Core | CCACGTTACTTAAAAGGCTGGTTGGAAGATGTGGTACAACTTTCGTTACGCCGCCCTAGCGTGCGTGCGAGCCGCCAACGTCC<br>AATCATTTCCTTGAATGAGCGCATTCTCGAATTCAATAAGCGTAACATTACAGCAATCATTGCTTATTACTTGCGTAAGTCGC<br>CGAGTGGATTGGATGTGGAGCGCGACCCGATTGAGTACGCCAAGTTTATGGAGCGCTATGCCGTGGGGTTATCGATTAAGACA<br>GAGGAGAAGTACTTCGACGGGTCCTACGAAACGCTGCGCAAGATCGCTTCATCGGTTTCCATCCCCATCTTAATGAATGACTT<br>TATCGTTAAGGAAAGTCAAATCGACGATGCATATAATCTGGGTGCGGATACTGTCCTGTTAATCGTTGAGATCCTTACAGAAC<br>GCGAGTTGGAGTCTTTGTTGGAGTACGCACGTTCCTATGGTATGGAGCCATTGATCCTTATCAACGACGAGAATGACTTGGAC<br>ATTGCGTTACGCATCGGTGCTCGCTTTATCACTATCTATTCTATGAACTTTGAGACCGGAGAGATCAATAAAGAGAACCAACG<br>TAAATTGATCAGTATGATTCCTAGTAACGTGGTGAAAGTGGCCCTTCTGGGGATTTCGGAGCGCAATGAAATTGAGGAGCTCC<br>GCAAGTTAGGTGTCAATGCATTTATGATTTCCAGCAGCCTGATGCGCAATCCGGAGAAGATCAAGGAGTTAATTGAAGGCAGC<br>CTTGAGCACCACCATCACCATCACTGA |
| RA95-Shell | CCGCGTTATTTAAAAGGATGGCTGGAAGATGTGGTTCAGTTGAGTCTTCGGCGACCCTCAGTTCATGCTTCCCGTCAGCGACC<br>AATTATTTCGCTCAATGAACGTATCCTGGAGTTTAATAAGCGCAATATTACGGCAATTATTGCCGTGTACGAGCGCAAGAGTC<br>CGAGCGGCCTCGATGTAGAACGAGATCCAATCGAGTATGCCAAGTATATGGAACCCTATGCGGTAGGTCTCAGCATAACCACT<br>GAGGAAAAGTATTTCAATGGCAGCTATGAGATGCTGCGCAAGATCGCCTCTTCCGTGTCCATACCCATATTAATGTCGGACTT<br>CATAGTTAAAGAAAGCCAAATTGATGATGCGTATAATCTTGGGGCGGACACAGTACTTCTGATTGTGAAAATATTAACCGAAC<br>GAGAACTGGAATCGTTATTGGAGTATGCCCGTGGATATGGGATGGAGCCTCTTATTTTGATAAACGATGAAAATGATCTTGAT<br>ATAGCTCTCCGCATTGGGGCACGCTTTATTGGGATAATGAGTCGTGATTTTGAAACTGGGGAAATCAATAAGGAAAACCAGCG<br>TAAGCTTATCAGCATGATCCCCAGTAATGTGGTGAAAGTTCCTAAATTAGATTTCTTTGAGCCCAATGAAATTGAGGAGCTCC<br>GTAAACTTGGTGTGAATGCGTTTCTTATCTCTTCAAGCCTGATGAGAAATCCGGAGAAAATAAAAGAACTGATCGAAGGTTCC<br>TTAGAGCATCACCATCATCATCACTGA |
| RA95-Core-Y51F | CCGCGCTATCTCAAAGGTTGGTTAGAGGACGTCGTTCAGCTTAGCCTGCGTCGTCCCAGTGTACGGGCTTCGCGCCAACGGCC<br>CATCATTAGCTTGAATGAGCGGATCCTCGAATTCAACAAGCGCAATATCACTGCTATAATTGCTTTCTATTTACGCAAAAGTC<br>CAAGTGGCTTGGATGTGGAGCGTGACCCTATCGAATATGCAAAGTTTATGGAGCGCTACGCGGTGGGACTTTCCATAAAGACA<br>GAGGAGAAATACTTTGACGGGTCTTACGAGACCCTGCGGAAGATAGCTAGCTCAGTCTCAATTCCTATATTGATGAATGATTT<br>TATTGTGAAGGAGTCACAGATAGACGATGCTTACAATCTTGGAGCTGATACAGTTTTACTGATAGTGGAGATTCTGACTGAGA<br>GAGAGTTGGAGAGCTTATTGGAGTACGCTCGTAGTTACGGCATGGAGCCTCTTATCTTAATTAATGACGAGAACGATCTTGAT<br>ATCGCCTTACGCATCGGGGCGCGTTTCATCACAATTTATAGTATGAATTTCGAGACCGGTGAGATCAACAAAGAGAATCAGAG<br>AAAATTAATCTCAATGATCCCGTCAAACGTGGTTAAGGTGGCCCTGCTCGGAATCTCTGAGAGAAACGAGATAGAGGAACTGC<br>GTAAATTGGGAGTGAACGCTTTCATGATCTCTAGCTCGCTTATGAGAAACCCTGAAAAGATCAAAGAGTTAATCGAAGGATCA<br>CTCGAGCACCACCACCACCACCACTGA |
| RA95-Core-N110S | CCCCGTTACTTGAAAGGTTGGCTGGAAGACGTGGTTCAATTAAGTCTCCGCCGGCCGTCGGTGCGCGCCTCTCGACAACGTCC<br>TATTATCTCCTTGAATGAAAGAATCCTGGAATTCAACAAGCGGAATATCACGGCAATAATAGCATACTACTTGCGCAAATCGC<br>CTAGCGGATTAGATGTGGAGCGTGACCCTATCGAGTACGCGAAGTTCATGGAACGTTATGCAGTTGGACTTTCAATTAAGACG<br>GAAGAAAAGTACTTTGACGGAAGCTACGAGACCCTGAGAAAGATCGCGTCTTCTGTCAGCATTCCAATTCTGATGTCCGACTT<br>TATTGTAAAGGAGTCTCAAATCGATGATGCGTACAACCTTGGCGCGGACACAGTCCTTCTTATTGTAGAAATTCTGACGGAAC<br>GAGAGTTAGAAAGTTTGCTTGAGTATGCACGAAGTTACGGCATGGAGCCACTCATCTTAATCAACGATGAGAATGATTTAGAC<br>ATCGCACTGCGCATAGGCGCCCGGTTTATCACTATTTACTCAATGAACTTCGAGACTGGTGAGATCAACAAAGAGAACCAACG<br>GAAACTGATATCAATGATTCCGTCGAATGTAGTAAAAGTCGCCCTTTTAGGGATTAGCGAAAGAAACGAGATTGAGGAGTTGC<br>GCAAGCTTGGGGTGAACGCCTTTATGATTTCTTCAAGCTTAATGCGGAACCCGGAGAAAATTAAAGAGCTGATAGAGGGTTCC<br>CTCGAGCACCACCACCACCACCACTGA |

| | |
|---|---|
| RA95-Core-Y180F | CCGCGCTACCTCAAAGGCTGGTTGGAAGACGTTGTACAACTGAGCTTACGGCGACCTAGTGTACGCGCTTCCCGTCAACGGCC<br>AATTATATCTCTGAACGAGCGCATCTTAGAGTTCAATAAGCGAAACATTACTGCAATTATCGCGTATTACTTGCGCAAAAGTC<br>CTTCTGGTCTTGATGTAGAGCGCGACCCTATTGAGTACGCGAAATTCATGGAGCGGTATGCAGTAGGTTTGAGCATCAAGACC<br>GAAGAGAAGTACTTCGACGGAAGCTACGAGACCTTACGCAAGATCGCTTCCTCAGTTAGTATACCGATTTTAATGAACGATTT<br>CATCGTCAAGGAGAGCCAAATCGACGATGCTTACAACCTGGGTGCCGACACCGTGCTGCTTATCGTAGAGATACTGACAGAGC<br>GGGAGTTGGAGTCTCTTTTAGAATACGCCCGTTCATACGGCATGGAACCTTTGATCCTTATCAACGACGAGAATGACCTGGAC<br>ATTGCATTACGCATTGGAGCCCGTTTCATAACTATCTTCAGTATGAATTTCGAGACGGGCGAGATTAACAAGGAGAATCAACG<br>CAAGCTTATTTCTATGATACCATCTAATGTAGTCAAAGTAGCATTGCTGGGTATTTCAGAACGCAATGAGATAGAAGAGTTGC<br>GCAAATTAGGCGTCAACGCATTCATGATAAGCTCCTCTTTAATGCGAAACCCGGAGAAAATCAAGGAGCTTATCGAGGGCTCT<br>CTCGAGCACCACCACCACCACCACTGA |

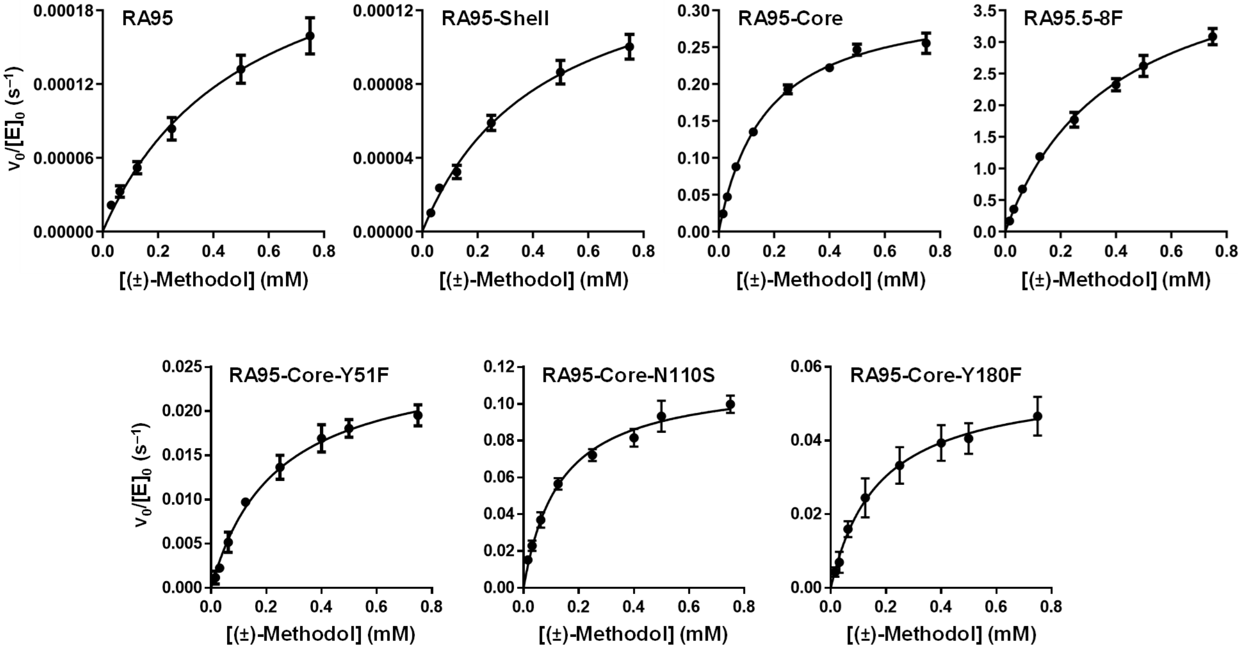


**Supplementary Figure 1. Steady-state kinetics.** Michaelis-Menten plots of normalized initial rates as a function of racemic methodol (4-hydroxy-4-(6-methoxy-2-naphthyl)-2-butanone) concentration are shown. Assays were carried out at 29 °C in 25 mM HEPES buffer (pH 7.5), 100 mM NaCl, 2.7 % acetonitrile. Product (6-methoxy-2-napthaldehyde) formation was monitored spectrophotometrically at 350 nm ($\varepsilon$ = 5,970 $M^{-1}$ $cm^{-1}$). Data represent the average of six or nine individual replicate measurements from two or three independent enzyme batches, with error bars indicating the SEM (n = 2 or 3 independent experiments, mean ± SEM in all cases). $k_{cat}$ and $K_M$ were determined by fitting the data to the Michaelis-Menten equation $v_0 = k_{cat}[E_0][S]/(K_M+[S])$.

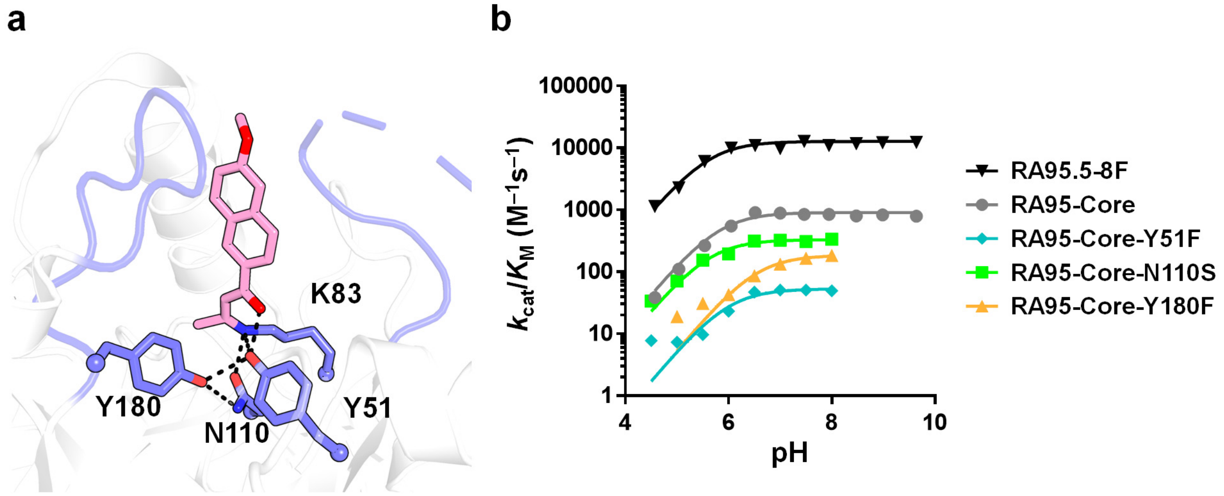


**Supplementary Figure 2. pH-rate profiles.** (a) Structure of the RA95.5-8F active site shows the catalytic tetrad and active site loops. (b) pH-rate profiles. Steady-state kinetic assays were carried out at 29 ºC in Britton-Robinson buffer at various pH with 2.7 % acetonitrile. Product (6-methoxy-2-napthaldehyde) formation was monitored spectrophotometrically at 350 nm (ε = 5,970 $M^{-1}$ $cm^{-1}$). Triplicate measurements were completed with varying concentrations of methodol (4-hydroxy-4-(6-methoxy-2-naphthyl)-2-butanone) at various pH. Initial reaction rates were fitted to the linear portion of the Michaelis-Menten model $v_0 = (k_{cat}/K_M)[E_0]$ by linear regression and $k_{cat}/K_M$ values were deduced from the slope. Data on the pH-rate profiles represent the $k_{cat}/K_M$ values determined at each pH. Error bars representing the errors of linear regression fitting to the linear portion of the Michaelis-Menten model are too small to be visible. These data were fitted to the following equation using nonlinear least squares regression: $(k_{cat}/K_M)_{obs} = (k_{cat}/K_M)_{max}/(1+10^{pK_{a1}-pH} + 10^{pK_{a2}-pH})$. The apparent $pK_a$ of the catalytic lysine ($pK_{a1}$) of each variant is presented in Table 1, with errors of nonlinear regression fitting provided.

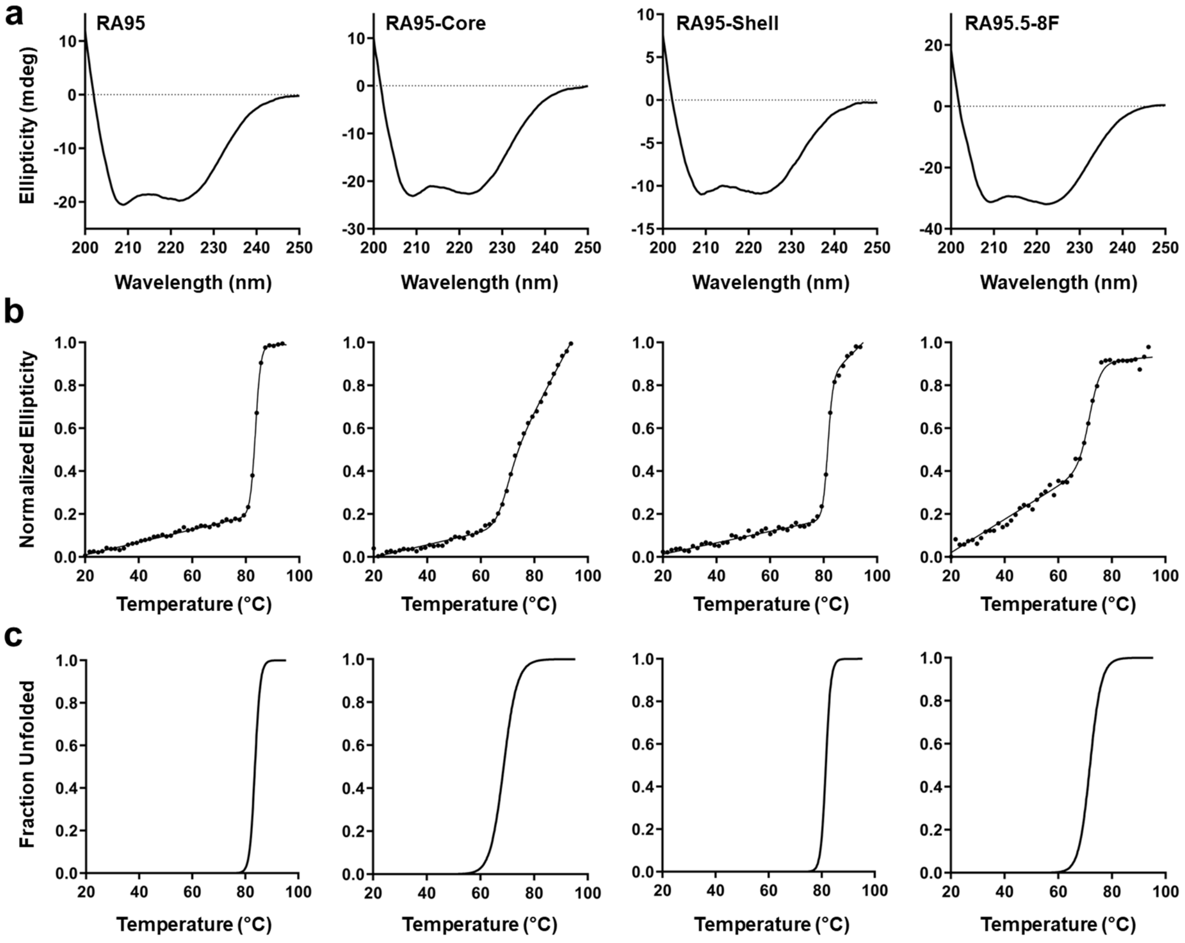


**Supplementary Figure 3. Circular dichroism and thermal denaturation assays for RA95 variants.** (a) Far-UV circular dichroism (CD) spectra. Scans were performed at 20 °C and sampled every 1 nm at a rate of 10 nm $min^{-1}$. Three scans were acquired and averaged for each sample. (b) Thermal denaturation monitored by CD at 222 nm. Samples were heated at a rate of 1 °C per minute, and ellipticity at 222 nm was measured every 0.2 °C ($n$ = 1). (c) Tm values were determined by fitting the data to a two-state transition model with correction for pre- and post-transition linear changes in ellipticity as a function of temperature (*45*) using nonlinear least-squares regression.

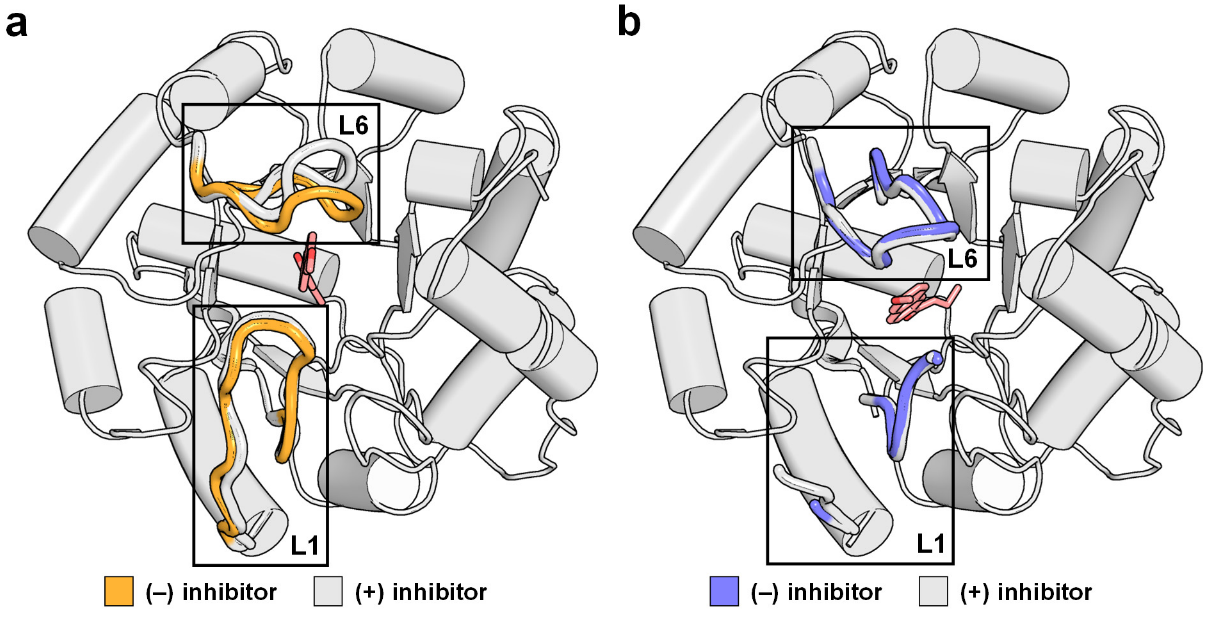


**Supplementary Figure 4. Active-site loops in RA95.5-8F are positioned for efficient substrate binding.** (a) Superposition of RA95 crystal structures with (white, PDB ID: 4A29) and without (orange, PDB ID: 9MYA) bound inhibitor (pink) show that loop L6 shifts position upon inhibitor binding. (b) By contrast, superposition of RA95.5-8F crystal structures with (white, PDB ID: 5AN7) and without (blue, PDB ID: 5AOU) bound inhibitor (pink) show that loop L6 does not shift upon inhibitor binding. Loops L1 and L6 are coloured in the bound structures. A representative retro-aldolase structure in grey is shown for the remainder of the protein. There is no electron density for loop L1 residues 58–61 and 58–63 in the 5AN7 and 5AOU structures, respectively, due to their high conformational heterogeneity.

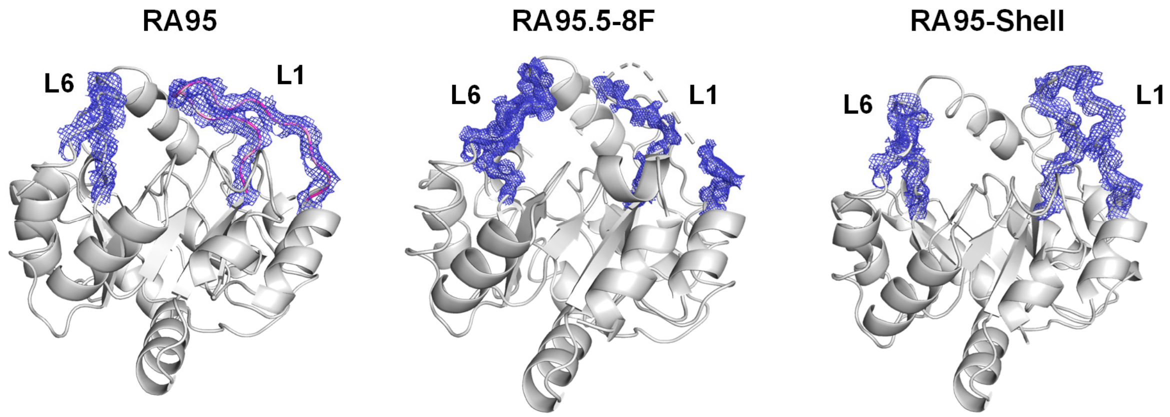


**Supplementary Figure 5. Electron density for loops L1 and L6.** 2mFo-DFc electron density maps contoured at 1 σ (blue mesh) for backbone atoms of loops L1 (residues 52–66) and L6 (residues 180–190) are shown for RA95 variants in their unbound forms. The missing density for loop L1 residues 58–63 in the crystal structure of RA95.5-8F is indicated by a dashed line.

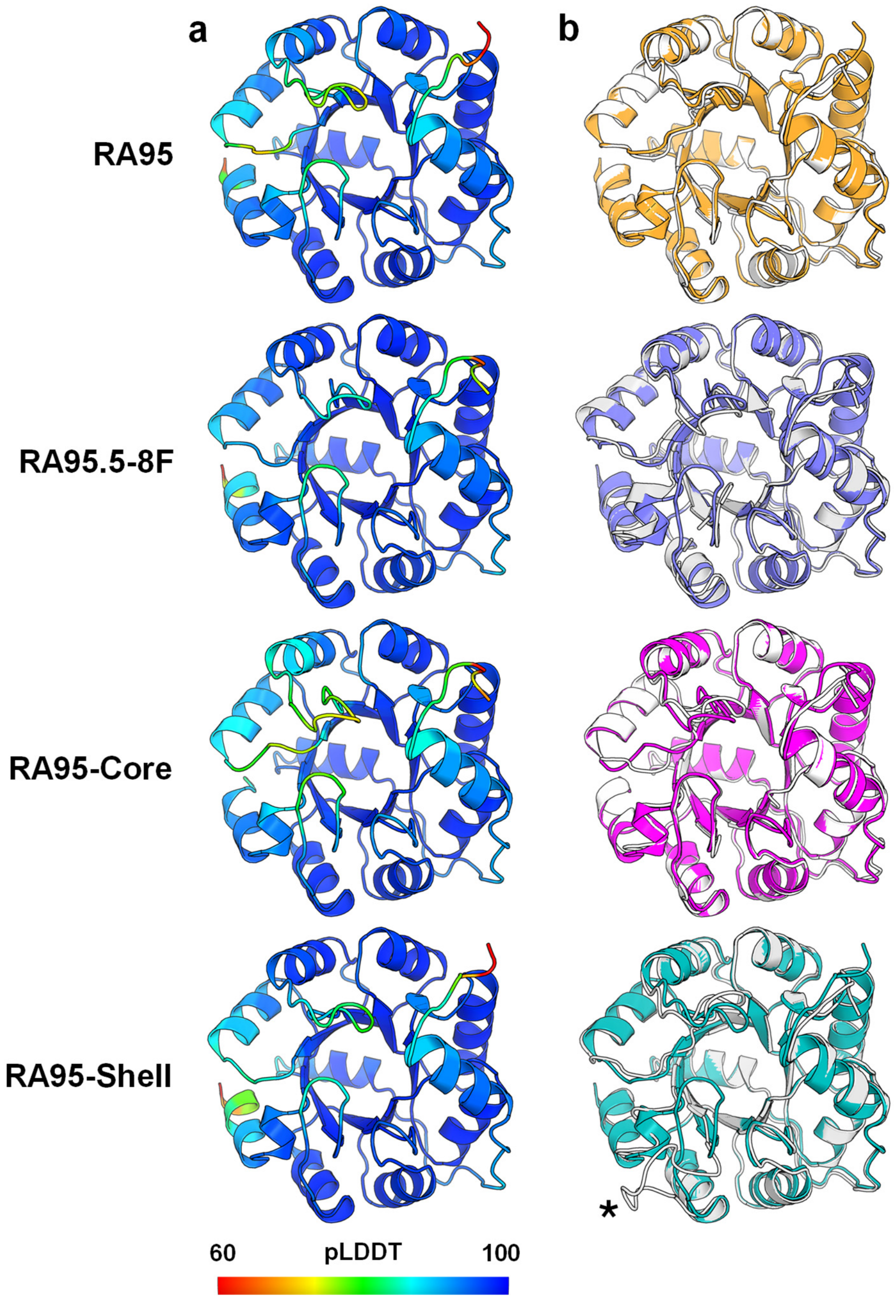


**Supplementary Figure 6. AlphaFold2 models of RA95 variants.** (a) AlphaFold2 models are coloured by pLDDT score. (b) Overlay of each model (coloured) with the corresponding unbound crystal structure (white). For RA95-Core, we used the RA95 crystal structure due to the absence of an RA95-Core structure. In all cases, AlphaFold2 predicts RA95-like active-site loops, unable to capture the open conformation of loop L1 seen in the RA95-Shell crystal structure (marked by an asterisk).

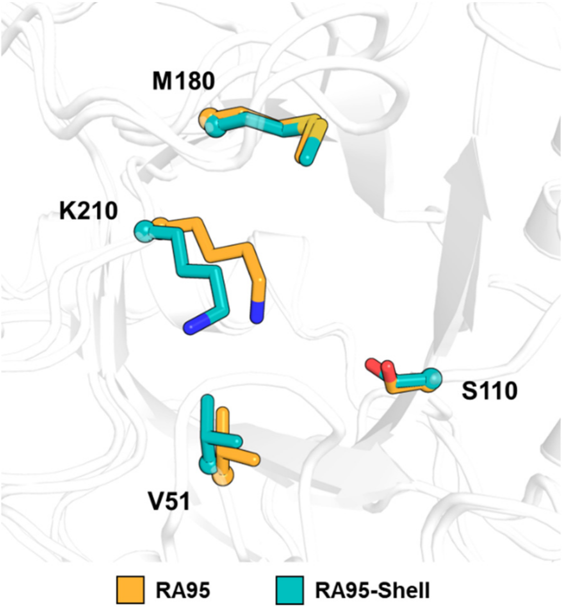


**Supplementary Figure 7. Distal mutations cause minimal changes to the rotameric configuration of active-site residues.** Superposition of active-site residues for unbound crystal structures of RA95 and RA95-Shell.

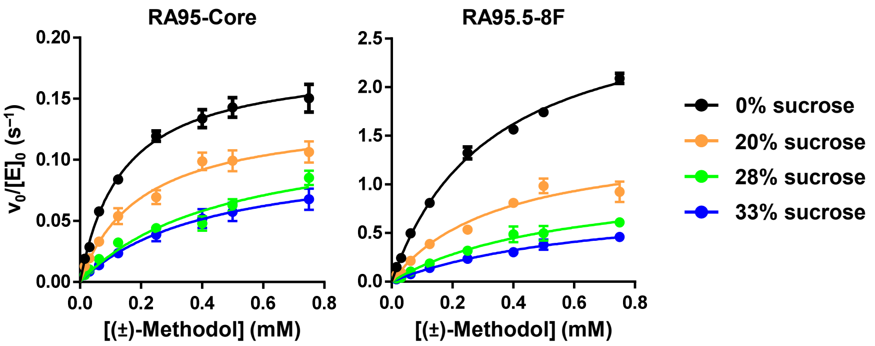


**Supplementary Figure 8. Steady-state kinetics at various viscosity.** Michaelis-Menten plots of normalized initial rates as a function of racemic methodol (4-hydroxy-4-(6-methoxy-2-naphthyl)-2-butanone) concentration are shown. Assays were carried out at 29 °C in 25 mM HEPES buffer (pH 7.5) supplemented with 100 mM NaCl, 2.7 % acetonitrile, and various sucrose concentrations. Product (6-methoxy-2-napthaldehyde) formation was monitored spectrophotometrically at 350 nm (ε = 5,970 $M^{-1}$ $cm^{-1}$). Data represent the average of six individual replicate measurements from two enzyme batches, with error bars indicating the SEM (n = 2 independent experiments, mean ± SEM in all cases). $k_{cat}$ and $K_M$ values were determined by fitting the data to the Michaelis-Menten equation $v_0 = k_{cat}[E_0][S]/(K_M+[S])$.

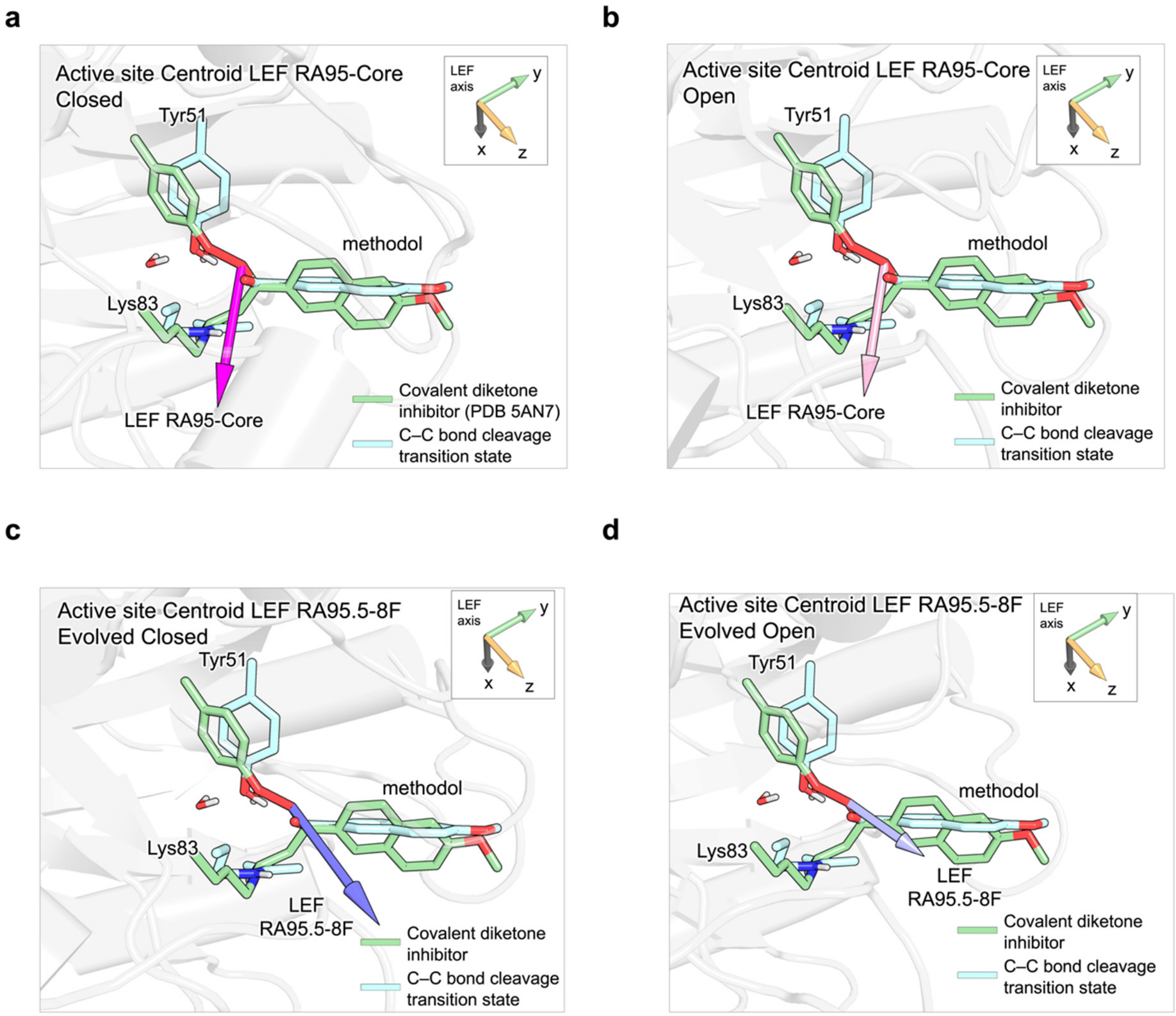


**Supplementary Figure 9. LEF of each variant (centroid structure from MD) with the theozyme C–C bond cleavage transition state aligned.** Active-site structures show the magnitude and direction of LEF vectors for each conformational state and variant: (a) RA95-Core Closed, (b) RA95-Core Open, (c) RA95.5-8F Closed and (d) RA95.5-8F Open. The theozyme transition state, including Lys83, Tyr51, and the methodol substrate, is shown in cyan sticks. Each enzyme centroid structure is aligned with the RA95.5-8F structure bound to a diketone inhibitor (PDB: 5AN7), with Lys83, Tyr51, and the inhibitor depicted as green sticks. The theozyme structure is aligned with active site residues and the inhibitor as described in the Methods section.

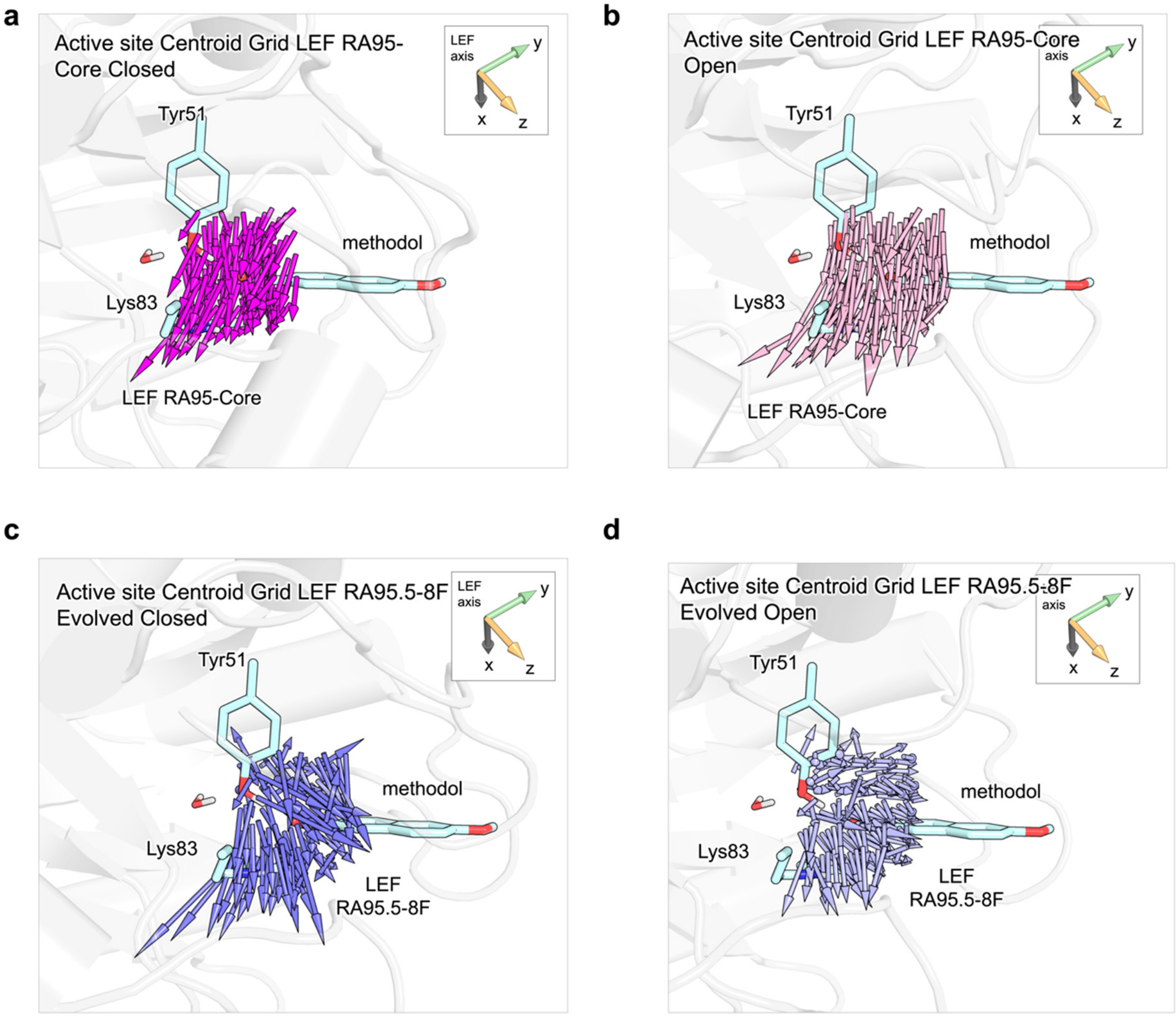


**Supplementary Figure 10. Calculated grid of LEFs in the active site of RA95 variants (centroid structures).** Active-site structures show the magnitude and direction of a grid of LEF vectors for each conformational state and variant: (a) RA95-Core Closed, (b) RA95-Core Open, (c) RA95.5-8F Closed, and (d) RA95.5-8F Open. The theozyme transition state, comprising the side chains of Lys83, Tyr51, and the methodol substrate, is shown in cyan sticks. Each enzyme centroid structure is aligned with the RA95.5-8F structure covalently bound to an inhibitor (PDB: 5AN7). The theozyme structure is aligned with active site residues and the inhibitor as described in the Methods section. A cubic box centered on the hydroxyl oxygen of the inhibitor was created, extending 2 Å in each direction along the X, Y, and Z axes, with a grid spacing of 1 Å, resulting in 125 points. The representative nature of the selected arbitrary point used for further analyses was proven by analyzing a grid of points in the active sites of the studied systems, confirming that this point effectively describes the trend of the LEF generated at each active site cavity.

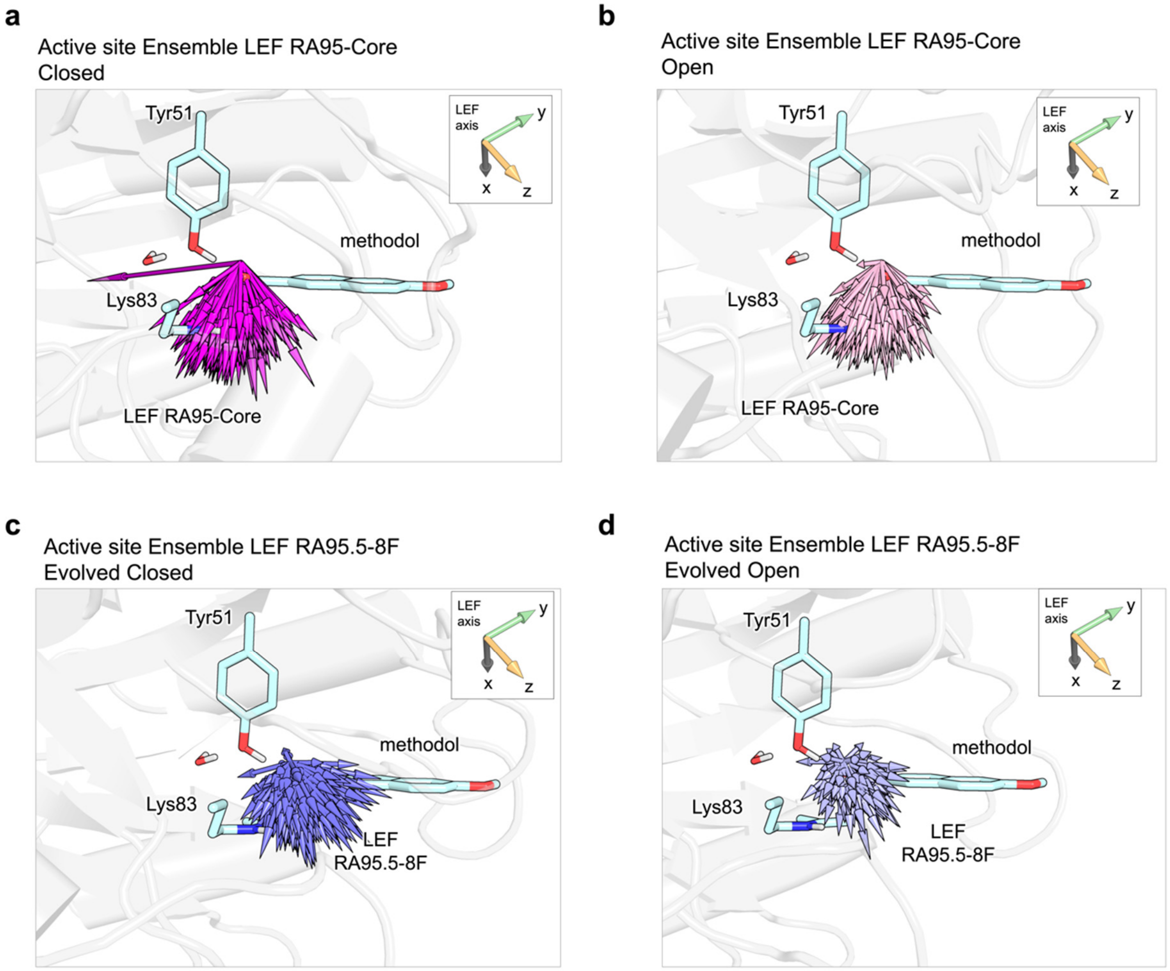


**Supplementary Figure 11. Calculated LEFs from the conformational ensemble of each variant derived from MD trajectories.** Active site structures show the magnitude and direction of superimposed LEF vectors calculated for each snapshot of the conformational ensemble from MD simulations for each conformational state and variant: (a) RA95-Core Closed, (b) RA95-Core Open, (c) RA95.5-8F Closed, and (d) RA95.5-8F Open. The theozyme transition state, including the side chains of Lys83, Tyr51, and the methanol substrate, is shown in cyan sticks. These results reveal that, despite some expected variations within the ensembles, the trends in both direction and magnitude of the calculated LEFs are consistently maintained and align closely with those obtained from the centroid structures.

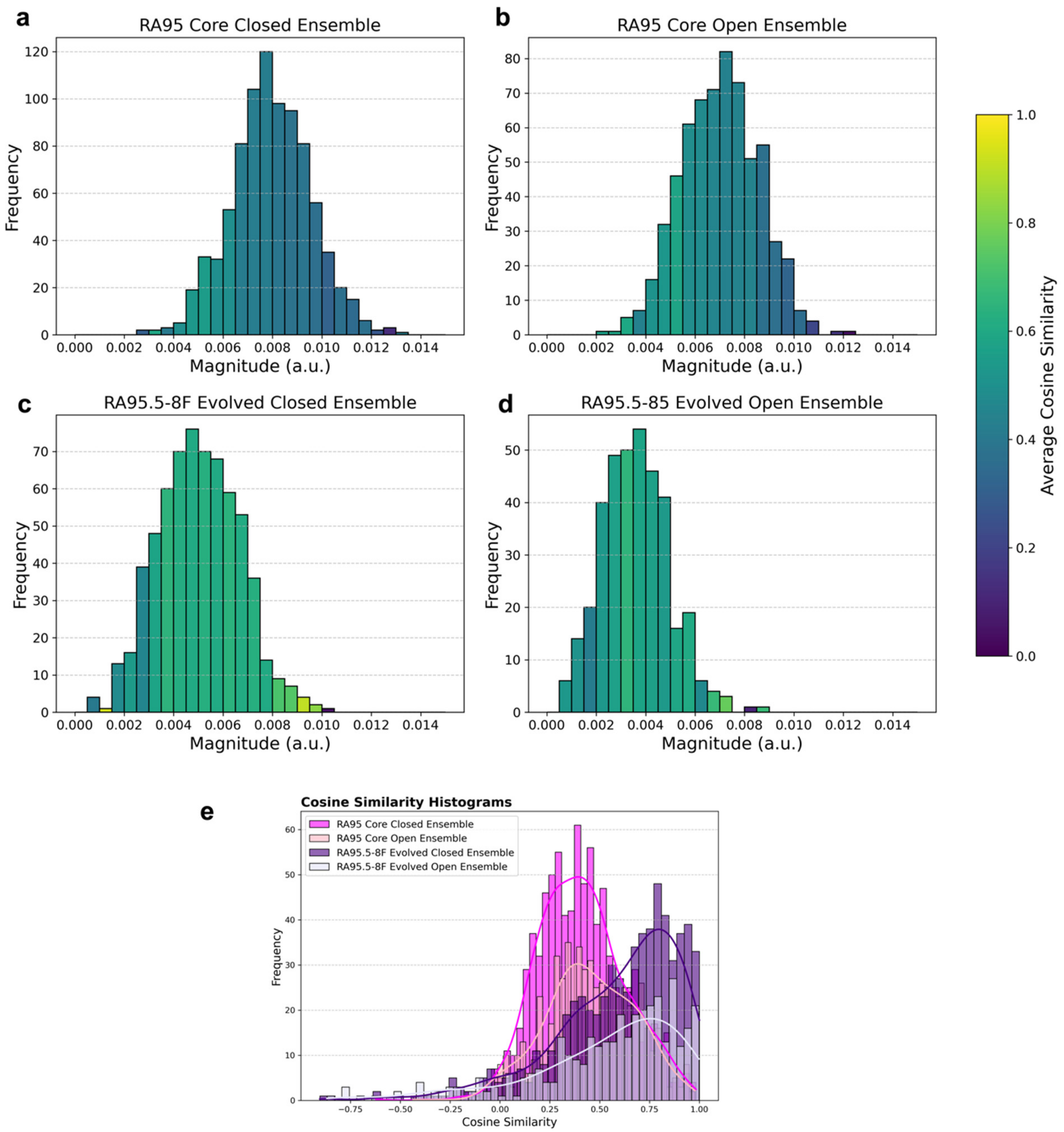


**Supplementary Figure 12. Analysis of LEFs calculated for the conformational ensemble of each variant derived from MD trajectories.** Histograms of active-site LEF ($\vec{F}$) magnitudes calculated for the conformational ensembles of: (a) RA95-Core Closed, (b) RA95-Core Open, (c) RA95.5-8F Closed, and (d) RA95.5-8F Open. The average cosine similarity (inner product space) between the LEF vectors in each bin, using the centroid RA95.5-8F Closed LEF as a reference, is represented by a color gradient. (e) Histogram of cosine similarity values (inner product) between LEF vectors for the conformational ensembles of each variant and state, using the centroid RA95.5-8F Closed LEF as a reference. These results demonstrate that the trends in LEF directions and magnitudes are consistent across the ensembles. Furthermore, the open and closed states of each variant exhibit similar LEFs.

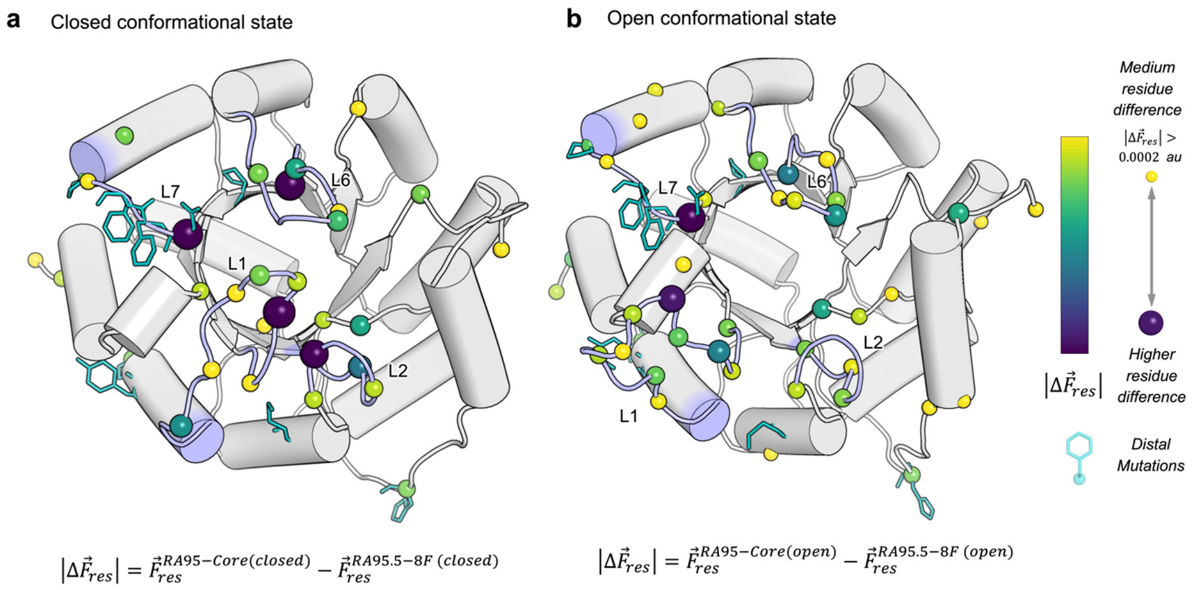


**Supplementary Figure 13. Residue-wise contribution to changes in the LEF.** Residue positions contributing the most to changes in the LEF are shown as coloured spheres for: (a) RA95-Core (Closed) to RA95.5-8F (Closed), and (b) RA95-Core (Open) to RA95.5-8F (Open). The size and colour of the spheres describe each residue's contribution to the LEF changes. The most significant changes in LEF originate from residues located on flexible loops (L1, L2, L6, L7), rather than directly from the distal mutation sites (shown as white spheres). The protein scaffold shown corresponds to RA95-Core (Closed) in (a) and RA95-Core (Open) in (b).
